# Global population structure and phase variation of serotype 12F *Streptococcus pneumoniae* following the introduction of pneumococcal conjugate vaccine

**DOI:** 10.64898/2026.04.03.714713

**Authors:** Thi N M Huynh, Alannah C King, Jason Chen Qixiang, Kathryn M Mulvihill, Hayley Demetriou, Kate C Mellor, Rebecca A Gladstone, Gemma G R Murray, Oliver Lorenz, Harry C H Hung, Teodora Mateeva, Sonu Shrestha, Sarah Kelly, Andrew J. Pollard, Shrijana Shrestha, John Lees, Samuel Horsfield, Feroze Ganaie, Sam Manna, Catherine Satzke, Lesley McGee, Chris Lok To Sham, David Goldblatt, Stephen D Bentley, Stephanie W Lo, The Global Pneumococcal Sequencing Consortium

**Author notes:** Corresponding Author Contact: Alannah C. King, Stephanie W. Lo.

## Abstract

**Background:** After the global deployment of pneumococcal conjugate vaccines (PCVs), serotype 12F has become the predominant serotype responsible for invasive pneumococcal disease (IPD) worldwide. As PCVs that include serotype 12F are gradually introduced, we aim to characterise the global population structure and genetic diversity of the 12F capsule locus using whole-genome sequencing. Capsule variants with vaccine evasion potential were further investigated by functional experiments.

**Methods:** A global collection of pneumococcal serotype 12F genomes (n=806) from 37 countries across six continents were included in this study. To characterise the serotype 12F population, Global Pneumococcal Sequence Cluster (GPSC), *in silico* serotype, and antimicrobial resistance profile were inferred from whole-genome data for each isolate. The capsule biosynthesis (*cps*) locus was analysed for gene content variations that could alter polysaccharide capsule production or structure, thereby influencing recognition by vaccine-induced antibodies. These isolates were further investigated by assessing their capsule production using immunofluorescence assays and its susceptibility to vaccine-elicited antibody killing by opsonophagocytosis assays.

**Findings:** The global increase in serotype 12F was driven by both distinct pneumococcal lineages across different continents, and a globally-disseminated and multidrug-resistant lineage GPSC26. We identified six capsule variants in nine isolates that had disruptive mutations in *cps* genes including *wze*, *wcil*, *wciJ* and *fnlA*. Most (6/9) of the disruptive mutations were a result of strand-slippage mutations. A convergent strand-slippage mutation disrupting the glycosyltransferase gene *wciJ* was identified in four isolates from distinct lineages and countries. Despite the truncation, three of four isolates with available Quellung typing results still identified them as 12F, indicating the production of the capsule. We then created a genetically engineered lab strain with *wciJ* knockout and complemented with *wciJ* containing the strand-slipppage mutation. The knockout strain did not produce any capsule. In contrast, the lab strain with *wciJ* containing the strand-slippage mutation produced a mixed population of encapsulated and non-encapsulated pneumococci, even within the same chain of pneumococcal cells. This observation indicated encapsulated subpopulation possesses a functional WciJ and rapidly reversible strand-slippage mutation during replication.

Opsonophagocytosis assays indicated that the clinical 12F strain with strand-slippage mutation in *wciJ* exhibited reduced susceptibility to vaccine-elicited serum killing, compared to a genetically closely related 12F clinical strain with an intact *wciJ*. However, substantial inter-individual antisera variation limits definitive interpretation.

**Interpretation:** Our work revealed the global rise of serotype 12F pneumococci has been driven by both regional-specific lineages, and a globally-disseminated and multidrug-resistant lineage GPSC26. We demonstrated that strand-slippage mutation is one of the major drivers of serotype 12F capsule variants and represents a novel mechanism enabling reversible on–off switching of capsule production. The ability to switch off capsule expression in a subpopulation may enable evasion of antibody-mediated killing but increase susceptibility to innate immune clearance.

**Funding:** Bill & Melinda Gates Foundation, Wellcome Sanger Institute, and the US Centers for Disease Control and Prevention.

## Introduction

*Streptococcus pneumoniae*, commonly known as the pneumococcus, can cause a spectrum of diseases, ranging from mild, localised infections such as otitis media to severe, invasive infections including sepsis and meningitis. Globally, the pneumococcus was estimated to have caused 829,000 deaths in 2019^1^ and ranks among the top three bacterial pathogens associated with the highest number of deaths attributable to antimicrobial resistance.^2^

Pneumococcal conjugate vaccines (PCVs) have proven effective in reducing pneumococcal disease worldwide. However, the increases in replacement of PCV serotypes with non-PCV serotypes remain a concern, particularly the global rise of serotype 12F following the introduction of PCVs. Serotype 12F has a high invasive disease potential,^3^ and has become the predominant serotype responsible for invasive pneumococcal disease (IPD) in multiple countries including Argentina,^4,5^ Canada,^6^ France,^7^ Israel,^8^ Italy,^9^ Japan,^10^ Korea,^11^ Germany,^12^, Netherlands,^13^ The Gambia,^14^ and Uruguay^15^ since the rollout of 7-valent PCV (PCV7), 10-valent PCV (PCV10), and 13-valent PCV (PCV13). Furthermore, serotype 12F has been linked to pneumococcal outbreaks among children,^16,17^ adults,^17,18^ individuals experiencing homelessness^19^ and prisoners.^20^ Multidrug resistance is also frequently observed in pneumococcal isolates of serotype 12F.^21–23^ The combination of high virulence, outbreak potential, and antimicrobial resistance makes the rise of serotype 12F a significant global health concern.

Serotype 12F is currently not included in the most widely used childhood pneumococcal vaccines, PCV10 (GSK or Serum Institute of India), PCV13 (Pfizer), and 15-valent PCV (PCV15, Merck), but is included in 20-valent PCV (PCV20, Pfizer) and higher-valency PCVs.^24^ PCV20 has now been licensed for paediatric use in the US and Europe since 2023/2024 and is increasingly being incorporated into routine childhood immunisation schedules. The 23-valent pneumococcal polysaccharide vaccine (PPV23, Merck), which included serotype 12F, was licensed in 1983 and has been used for older adults and those with risk conditions. PPV23 shows effective protection against serotype12F IPD cases among older adults.^25^, Instances of vaccine failure to protect against 12F IPD were rare, with only one reported case of an adult developing 12F IPD, which occurred two years after receiving PPV23 in Japan.^26^

As PCVs that include serotype 12F are gradually being introduced, we aim to characterise the global population structure and genetic diversity of 12F pneumococci through whole–genome sequencing. Additionally, capsule variants were assessed for their capsule production using immunofluorescence assay and their susceptibility to vaccine-elicited antibody killing by opsonophagocytosis assay.

## Research in context

### Evidence before this study

We searched PubMed using the terms “*Streptococcus pneumoniae*” AND “12F” for the papers published in English between January 1, 2020 and July 15, 2025. We searched for studies which reported the changes in pneumococcal serotype or/and population before and after the introduction of PCVs in countries or regions. After reviewing 87 articles, 55 met the inclusion criteria for this research topic.

Out of 55 articles, 5 were meta-analysis studies conducted in high-income countries (n=3) and globally (n=2); the impact of PCV10 was evaluated in 5 studies in Belgium, Ethiopia, Netherlands and Nigeria, PCV13 was assessed in 44 studies in Argentina, Brazil, Canada, Democratic Republic of the Congo, France, Ghana, Israel, Japan, Scandinavian countries, South Africa, Spain, UK and US. Additionally, PCV15 was examined in 1 study in Greece. Most studies reported declines in IPD cases in children and adults (n=19), adult (n=17), or children (n=14); There were also two studies on non-invasive pneumococcal disease cases in children and three studies on asymptomatic colonisation in children.

Of the 50 IPD studies reviewed, 33 reported an increase in serotype 12F or identified it as among the top five serotypes. This trend was noted in children and adults in Argentina, Brazil, Canada, Israel, Japan, Spain, and the UK; among children in France, Nigeria, South Africa, and the US; and among adults in the Netherlands, and Scandinavian countries. Following a prior increase, a decline in serotype 12F-associated IPD was observed in Canada, the Netherlands, and the UK during and after the COVID-19 pandemic in 2020 and also observed in Brazil six to eight years after the introduction of PCV10. In contrast, none of the studies on non-invasive pneumococcal disease or carriage studies identified serotype 12F as a common serotype, except for Democratic Republic of Congo, reinforcing the observation that serotype 12F is primarily associated with invasive disease and exhibits high invasive potential.

Only seven studies characterised the serotype 12F strains using multi-locus sequence typing and/or genomic lineage analysis. GPSC32 was identified in Canada, US, and Spain, GPSC26 in Spain and Nigeria, GPSC55 in Israel and Spain. However, no study has reported on the global population structure of serotype 12F or examined the serotype 12F capsular region in detail.

### Added value of this study

By utilising a global collection of *S. pneumoniae* serotype 12F isolates from 37 countries, we have enhanced our understanding of this highly invasive serotype, which has been responsible for multiple outbreaks and has contributed to the global serotype replacement following the introduction of PCV7, PCV10 and PCV13. This study characterised the global population structure of serotype 12F *S. pneumoniae*, its antimicrobial resistance profiles, and explored the genetic diversity of serotype 12F capsule locus. We showed strand slippage mutation is one of the major mechanisms for 12F capsule diversification, a novel mechanism for capsule phase variation that could occur rapidly during replication, and such heterogeneous production of capsules in a single strain may reduce susceptibility to vaccine-elicited antibody-mediated killing.

### Implications of all the available evidence

The increase of serotype 12F has been driven by distinct regional lineages, and a globally-disseminated and multidrug-resistant lineage GPSC26. The time-resolved phylogeny in this study showed that GPSC26 is likely to have emerged around 1943 and expanded in 1988, a time when PPV23 was not widely utilised worldwide and prior to the introduction of PCVs. Our study showed that GPSC26 was mainly found in Africa and Asia. At the time writing, our parallel studies showed increasing detection of GPSC26 in Europe and South America.

We identified previously unrecognised genetic diversity within the capsule biosynthesis locus of serotype 12F and uncovered a novel mechanism - strand slippage mutation in glycosyltransferase - to enable a reversible on-off switching capsule production. Such heterogeneity in capsule expression within a population may enhance pneumococcal survival by adherence, pathogenesis^27^ and reducing susceptibility to vaccine-elicited, serotype-specific immune responses, as non-encapsulated subpopulations are inherently more resistant to vaccine-elicited antibody killing.

## Methods

### Study design

A global collection of serotype 12F pneumococcal genomes (n=834) was included in this study from the Global Pneumococcal Sequencing (GPS) database (n=624, last accessed on 25^th^ March 2022)^28^ and from five previous studies with genome data not in the GPS database (n=210)^10,17,29–31^ (Table S1). The genomes underwent quality control and bioinformatic analyses and were collated with the epidemiological data collected. To characterise the pneumococcal lineages contributing to the increase in serotype 12F, we grouped isolates into Global Pneumococcal Sequencing Clusters (GPSCs) and contextualised the 12F lineage of interest by incorporating additional genomes of this lineage from the GPS database, irrespective of serotype. We examined the capsule biosynthesis (*cps*) locus which encodes for the vaccine antigen for variants, focusing on differences in gene content contributed by premature stop codons due to frameshift or missense mutation. These mutations could potentially change the polysaccharide capsule production or structure, affecting the response of vaccine-elicited antibodies to the antigen and leading to vaccine evasion.

### Genome sequencing and analysis

The pneumococcal genomes from the GPS database in this study were whole-genome sequenced by Illumina HiSeq or NovaSeq at the Wellcome Sanger Institute (Hinxton, UK) while pneumococcal genomes from the previous five studies^10,17,29–31^ were sequenced by MiSeq sequencing platform. Velvet (v1.2.10)^32^ and Spades (v3.1)^33^ were used to *de novo* assemble raw read data from HiSeq and MiSeq sequencers, respectively. The raw reads and assemblies were subjected to quality control as previously described.^34^ The accession numbers, metadata and in *silico* typing output were included in Supplementary Data S1.

For each genome, *in silico* serotype was inferred using SeroBA with default parameters (v1.0.0). ^35^ Population Partitioning Using Nucleotide K-mers (PopPUNK) (v2.4.0) was used to cluster isolates into GPSC using the reference GPSC database v6.^36,37^ Sequence types (STs) of the genomes were inferred by multilocus sequence typing (MLST) (v.2.9)^38^. Resistance to penicillin was predicted based on PBP1a, PBP2X and PBP2b amino acid patterns using a machine learning model.39,40 The predicted MIC values were interpreted using CLSI M100 guideline. Penicillin resistance was defined as MIC of ≥0.12 µg/ml, according to meningitis breakpoints. Resistance to chloramphenicol (presence of *cat*), cotrimoxazole [I100L mutation in *folA* and indel between amino acids 56-67 in *folP*], erythromycin (presence of *erm(B)* or *mef(A)*), tetracycline (presence of *tet*(M), *tet*(O) or *tet*(S/M)), and vancomycin (presence of *van*(A), *van*(B), *van*(C), *van*(D), *van*(E) or *van*(G)), were also predicted for each genome using the Antimicrobial resistance (AMR) pipeline developed by the US CDC.^39,41^ Isolates were defined as multidrug resistant (MDR) when predicted to be non-susceptible to ≥3 antimicrobial classes.

### Phylogenetic analysis

We performed phylogenetic analysis on all serotype 12F genomes by constructing a maximum likelihood tree using FastTree (v2.1.10) with a general time reversible substitution model.^42^ Phylogenies were built based on the single nucleotide polymorphisms (SNPs) extracted from an alignment generated by mapping reads to a reference genome *S. pneumoniae* ATCC 700669 [National Center for Biotechnology Information (NCBI) accession number FM211187] using SMALT (v0.7.4) with default settings.^43^ This reference genome has been used throughout the whole GPS project. The phylogenetic trees were then overlaid with epidemiological data and the *in silico* output, and visualised in Microreact, available at https://microreact.org/project/gps-global-12f-pneumococci.

### Evolutionary and spatiotemporal analysis

Sequence reads of GPSC26 isolates were mapped to the GPSC26 reference genome (Genbank accession number LS483450.1) using Burrows Wheeler Aligner (version 0.7.17-r1188).^44^ Recombination was detected, removed, and a recombination-free phylogeny was constructed with RAxML using Gubbins (version 3.2.1).^45^ The phylogeny was visualised in Microreact, available at https://microreact.org/project/gps-global-gpsc26.

To construct a time-resolved phylogeny, we first investigated the presence of temporal signals by a linear regression of root-to-tip distances against year of collection using TempEst (v1.5.3) ,^46^ and excluded genomes located on long branches that distorted the temporal signal. Then, Bayesian phylogenetic analysis on a subset of serotype 12F isolates with temporal signal was used to generate a time-resolved phylogeny and provide estimates of the median effective population size over time with a 95% highest posterior density to detect any exponential increase in population size using BEAST Bayesian skyline model (version 1.10.4).^47^ Three independent Markov Chain Monte Carlo (MCMC) runs of 100 million generations each were performed, sampling every 10,000 states. All parameters had effective sample sizes (ESS) > 200, indicating convergence. A relaxed molecular clock was applied, with GTR+ Gamma as the nucleotide substitution model. The time-resolved phylogeny was visualised in Microreact, available at https://microreact.org/project/gps-gpsc26-beast.

### Genetic analysis of serotype 12F capsular encoding region (*cps*)

To examine the genetic variation of the vaccine target, we extracted the *cps* sequences from the assemblies of all serotype 12F pneumococcal genomes. We then used an in-house script to identify differences in gene content and the presence of pre-mature stop codons in each *cps* gene.^48^ Detected premature stop codons were validated by manual inspection of raw read alignments against the assembled *cps* sequences.

In addition, a phylogeny of the serotype 12F *cps* was built by mapping raw reads of each genome to a 12F *cps* reference sequence (NCBI accession number CR931660) using Burrows Wheeler Aligner (BWA) (v 0.7.17),^44^ and a maximum likelihood tree was constructed using FastTree (v2.1.10) with a general time reversible substitution model.^42^ The 12F *cps* phylogeny was then visualised in Microreact at https://microreact.org/project/gps-12f-cps.

### Immunofluorescence assay of capsule production

Pneumococcal strain D39W and its derivative constructs are summarised in Table S2 while primers and DNA templates for constructing the strains are detailed in Table S3. PCR amplicons were amplified using Phusion DNA polymerase (NEB, M0530S) with the appropriate reagents as per the manufacturer’s instructions. Amplicons were confirmed via gel electrophoresis and purified using the QIAquick PCR purification kit (Qiagen, 28106). PCR amplicons were assembled using Gibson assembly^49^ to generate genetic cassettes for strain construction. These cassettes were introduced to pneumococcus by inducing natural transformation. Briefly, strains were cultured in BHI broth to an OD600 of 0.1 to 0.3. Cultures were then normalised to an OD600 of 0.03 before adding competence-stimulating peptide-1 (CSP-1) to induce competence. After which, the genetic cassette was added, and the mixture was incubated for another 90 min at 37°C with 5% CO_2_. Transformation cultures were plated on blood agar supplemented with the appropriate antibiotics. Strains were then validated by PCR using 2X PowerPol polymerase (ABclonal, RK20718) and primers flanking the cassette. Amplicons were verified by Nanopore sequencing.

Heat-killed Cells at the density of OD600 0.2-0.4 were then resuspended in 100 µl of cross-absorbed anti-CPS12 sera (SSI Diagnostica) at a 1:600 dilution and incubated on ice for 5 minutes. The anti-CPS12 sera can detect all serotypes within serogroup 12. Afterwards, cells were washed twice with 1 ml of 1X PBS and resuspended in 100 µl of 1X PBS. Anti-rabbit Alexa Fluor 488 secondary antibody (A11034) that emits green fluorescence was added to the mixture at 1:100 dilution and incubated on ice for 5 minutes. Cells were washed one more time with 1 ml of 1X PBS before being resuspended in 100 µl of 1X PBS and visualized with an Eclipse Ti2 Microscope (Nikon). The experiments were carried out in at least two biological replicates and all the images were collated in figure S6-10.

### Opsonophagocytosis Assays

The opsonophagocytosis assay (OPA) was performed in the World Health Organisation reference laboratory for pneumococcal serology reference laboratory (PSRL) at the Great Ormond Street Institute of Child Health, University College London. The 12F pneumococcal isolates for testing were incorporated in the assay and run together with a validated 12F OPA reference strain DS4009-06 (GenBank: JAXIIY000000000.1, GPSC32, ST220) developed in the PSRL department. Post vaccination sera were collected from healthy adults 4-6 weeks after immunisation using PPV23 in a 2004 clinical trial.^50^

The OPA methodology was previously described^51^ and was adapted to test a single pneumococcal strain at a time. In brief, the 10 patient sera were diluted three-fold from 1:1 to 1:8 and then incubated with the known concentration of bacterial strain at room temperature (19 - 25°C) on a shaker (700 rpm) for 30 minutes. Then, differentiated HL60 cells and baby rabbit complement (BRC) were added and incubated at 37°C on a shaker for 45 minutes. The mixture was then subject to eight-fold dilutions and each dilution was spotted onto THY agar plates, coated with triphenyl tetrazolium chloride (TTC) incubated for 16-18 hours at 37°C. Bacterial colonies on the agar plates were counted using an automated colony counter (ProtoCol 3, Synbiosis Ltd.). OPA index (OI) was determined by the sera dilution of titer at which 50% of the bacteria were killed. The experiment included the 12F OPA reference strain with known OPA titer as control. A value of 5.5 was assigned to samples with an OI titre below the lower limit of detection.

### Statistical tests

To determine if differences in proportions between groups were significant, a two-sided Fisher’s exact test was used. To compare OPA index, geometric means and 95% confidence intervals (CIs) for each group were calculated, and the difference was detected using one-way analysis of variance (ANOVA) and Turkey’s Honest significant difference test was used for pairwise post hoc comparisons. Multiple testing correction was carried out using the Benjamini-Hochberg false discovery rate of 5%. Two-sided p values of less than 0.05 were considered significant. The statistical tests were performed in R, version 3.6.0 or above.

## Results

### Bacterial collection

Among 834 serotype 12F pneumococcal genomes, 806 (96.6%) from 37 countries across six continents met the quality control criteria. The collection included disease (87.3%; n=704) and carriage isolates (7.1%; n=57), while 5.6% (n=45) of samples were of unknown clinical manifestation. The samples were collected from individuals aged between 0 and 98 years old between 1990 and 2019.

### Population structure and antimicrobial resistance

The isolates were clustered into nine GPSC lineages: GPSC26 (CC989; n=359; 44.5%), GPSC32 (CC218; n=166; 20.6%), GPSC334 (CC4846 and CC1527; n=101; 12.5%), GPSC55 (CC3524 and ST3776; n=101; 12.5%), GPSC56 (CC8672; n=64; 8.0%), GPSC212 (ST6202; n=12; 1.5%) and one isolate each from lineages GPSC575, GPSC289, and GPSC648 (totaling 0.4%). The lineages driving the increase in serotype 12F varied across countries and continents (Figure 1A). Overall, 94.0% of GPSC32 samples originated from the Americas, all GPSC56 samples were from South Africa, 90.1% of GPSC55 samples were from Israel, and 93.1% of GPSC334 samples were from Japan. In contrast, GPSC26 was identified in 26 of the 37 countries, predominantly in Africa, to a lesser extent Asia and Europe (Figure 1B).

**Figure 1.**
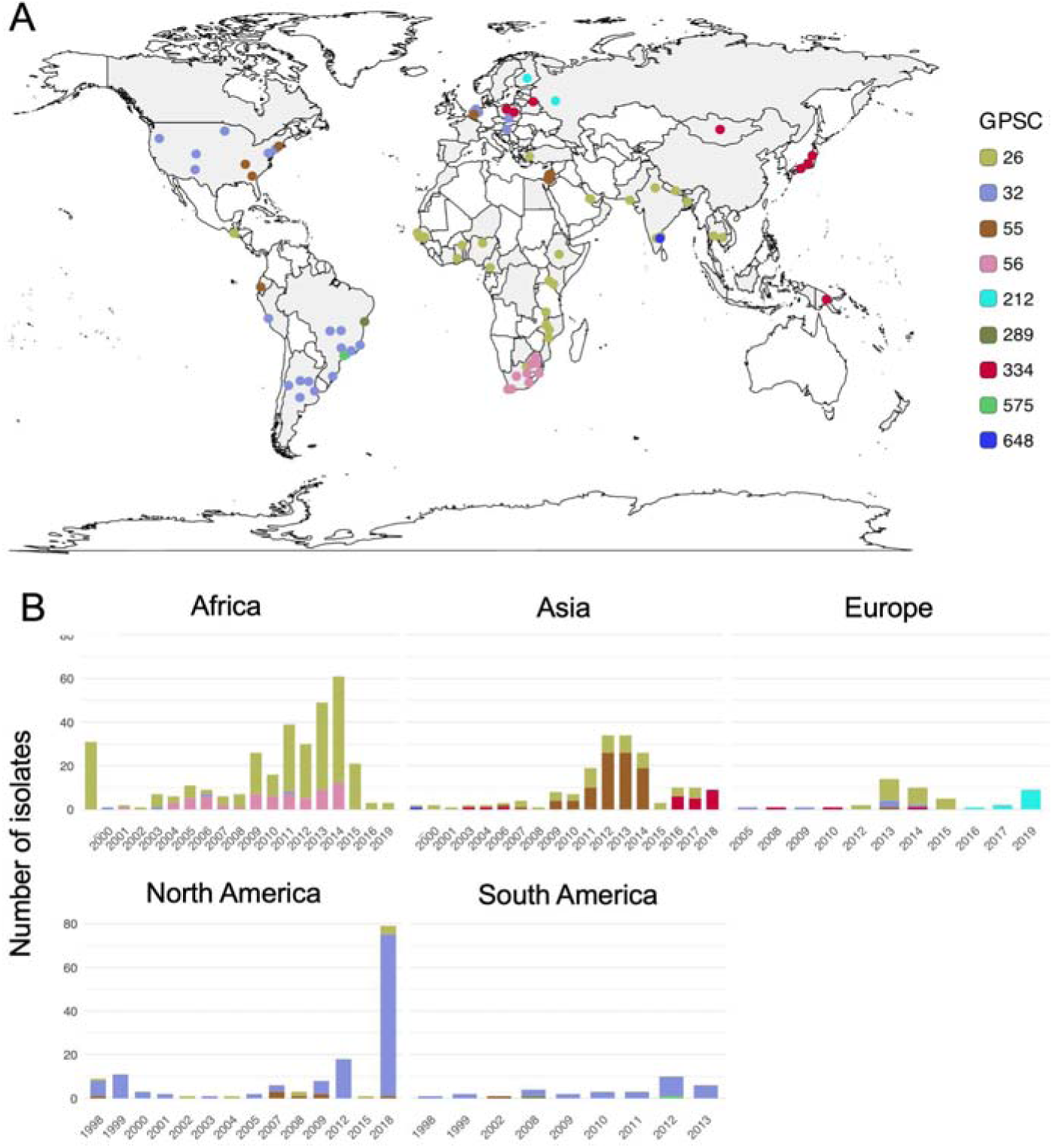
Global population structure of serotype *12F Streptococcus pneumoniae* (n=806) (A) Global distribution of serotype 12F pneumococcal lineages; (B) Proportion of serotype 12F pneumococcal lineages over study period by continents; Oceania (n=1) and isolates with unknown continent (n=1) and year of collection (n=108) are excluded.

Of the nine serotype 12F lineages, GPSC26 and GPSC334 were identified as MDR lineages (Figure 2 and Table S4). MDR was observed in 92% (n=330/359) of GPSC26 isolates, with 91.7% (n=329/359) resistant to tetracycline, 77.7% (n=279/359) resistant to chloramphenicol, and 100.0% (n=359/359) intermediate or fully resistant to cotrimoxazole. Additionally, 92.5% (n=332/359) of GPSC26 isolates were predicted to remain susceptible to penicillin, with only 27 isolates exhibiting resistance. The global phylogenetic analysis of GPSC26 identified three clusters of penicillin-resistant isolates, each exhibiting distinct *pbp* profiles. These findings indicated independent acquisitions of penicillin resistance, followed by clonal expansions in the southern region of Africa, the Middle East and Eastern Europe, and Nigeria (Figure S1). Penicillin resistance was also identified in a single isolate each from Bangladesh and Cambodia.

**Figure 2.**
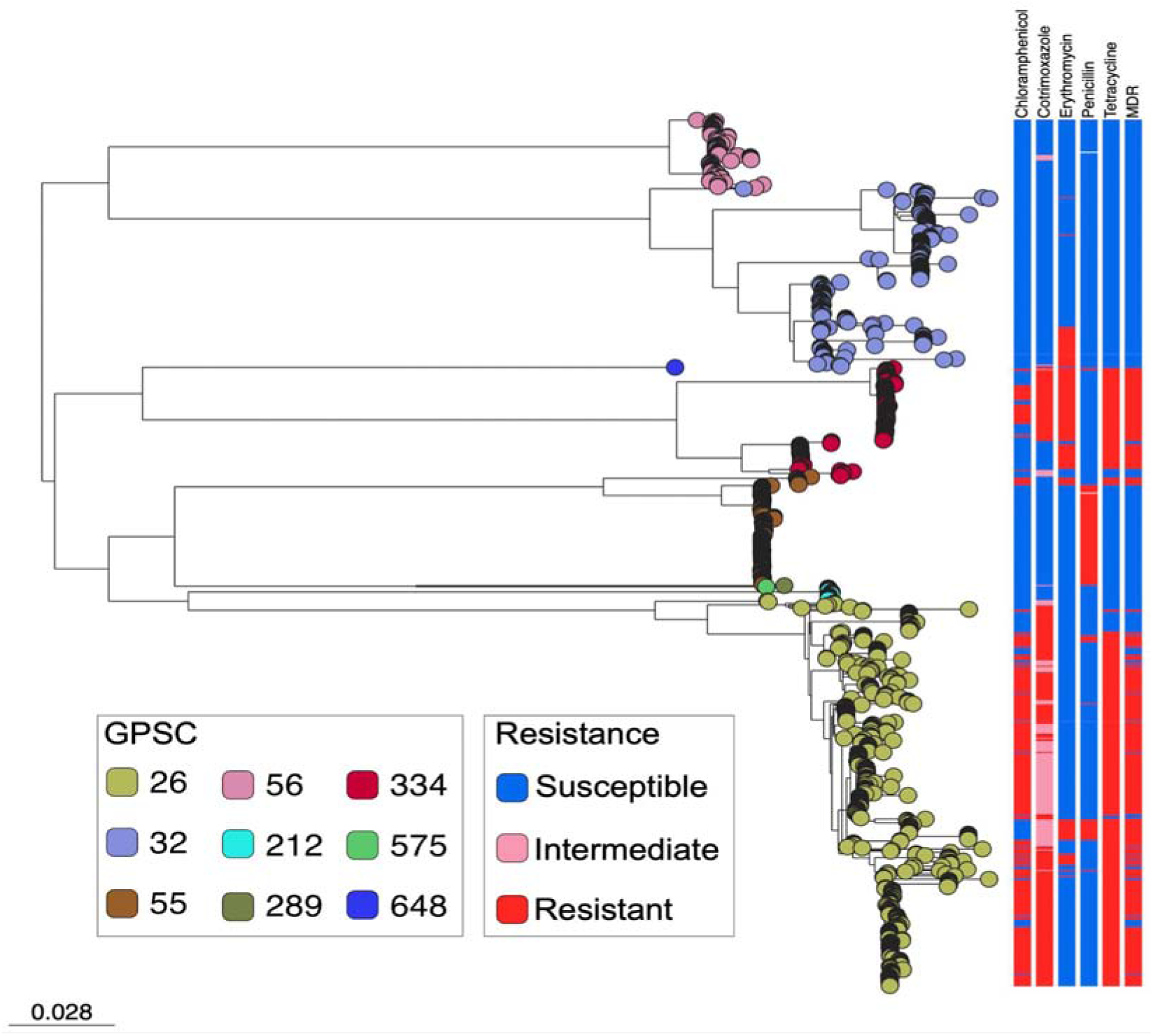
Phylogeny of serotype 12F pneumococcal isolates. (n=806) based on single-nucleotide polymorphisms (SNPs) across the genome, overlaid with antimicrobial resistance profiles. The scale bar refers to a genetic distance of 0.028 nucleotide substitutions per site. Penicillin resistance was based on the predicted MIC at ≥0.12 µg/ml, according to meningitis breakpoints.

The majority (90.1%, n=91/101) of GPSC334 isolates were MDR, with 90.1% (n=91/101) resistant to erythromycin, 93.1% (n=94/101) resistant to tetracycline, 74.3% (n=74/101) non-susceptible to cotrimoxazole, and 36.6% (37/101) resistant to chloramphenicol. Additionally, 63.4% (n=64/101) of the GPSC334 isolates had a susceptible penicillin *pbp* profile *pbp1A-pbp2B-pbp2X* pattern of 37-278-375. Only two GPSC334 isolates were resistant to penicillin due to the acquisition of *pbp2X* variants (37-278-498 and 37-278-1049) individually. Notably, the isolate with *pbp* pattern 37-278-1049 was non-susceptible to cefuroxime.

Among the 12F pneumococcal lineages, GPSC55 was the only lineage where nearly all isolates (91.1%, n=92/101) were resistant to penicillin. A cluster of seven GPSC55 isolates were found to be penicillin-susceptible but MDR to chloramphenicol, erythromycin and tetracycline. These isolates were primarily detected in the USA (n=6/7), with one isolate identified from Ecuador (n=1/7). No MDR isolates were observed in GPSC32, GPSC56, GPSC212, GPSC289, GPSC575, and GPSC648.

### Globally-disseminating serotype 12F pneumococcal lineage GPSC26

We conducted a further investigation into the globally-disseminating serotype 12F lineage GPSC26 using an international collection of 436 GPSC26 isolates, irrespective of their serotypes. This lineage was identified in 28 countries, with the following distribution: Africa (n=298), Asia (n=98), Europe (n=25), North America (n=10), South America (n=1), Oceania (n=1) and one isolate of unknown origin. The majority of GPSC26 isolates expressed serotype 12F (n=359), along with genetically and structurally related serotypes 46 (n=55), 12B (n=9), and 12A (n=3). Few isolates expressed serotypes included serotype 9L (n=5), and one isolate each of serotype 9V, 15A, 19F, and 40.

The global phylogeny of GPSC26 (Figure S2) revealed multiple independent acquisitions of serotype 46 *cps*, which were followed by local expansions in several countries including The Gambia, Kenya, India, Malawi, Bangladesh, Israel, and Nigeria (Figure S3). Similar patterns of *cps* acquisitions and subsequent expansion were also noted for serotype 12A in Bangladesh and serotype 9L in Ethiopia. Other serotypes emerged sporadically, with no indication of clonal expansion, suggesting that the genetic background of GPSC26 may be less compatible with these other serotypes.

A time-resolved phylogeny was constructed using 398 GPSC26 genomes, after excluding 38 outlier genomes identified through the linear regression of root-to-tip distance against year of collection (Figure S4). These highly divergent GPSC26 isolates were excluded to avoid biasing the molecular clock and coalescent inferences. This exclusion improved the temporal signal, increasing the R^2^ value from 0.04 to 0.31, which enabled convergence in the BEAST analysis, indicated by the effective sample size (ESS) >200. GPSC26 is estimated to have emerged around 1943 (with a 95% highest posterior density 1916-1967), and it is predicted to have experienced an exponential increase in effective population size around 1988 (Figure S5). Although PPV23 was licensed and used in the US since 1983, it was not widely used in Africa and Asia until the 1990s. Therefore, the predicted expansion was probably not linked to any vaccine introduction.

By comparing the phylogenies constructed from SNPs across the genome and *cps* locus, we identified three potential capsular switching events (Figure S6). GPSC55 may have served as a 12F *cps* donor for GPSC289 (n=1), while GPSC26 may have been the donor for GPSC212 (n=12) and GPSC648 (n=1), as indicated by their high nucleotide similarity in *cps* and dissimilarity in the genetic backgrounds.

### Genetic variants of the serotype 12F capsule and diversification mechanism

Among the 806 serotype 12F genomes, an intact *cps* locus was extracted from 804 genomes. We identified six *cps* variants in nine isolates that had disruptive mutations in initial transferase WciL (n=1), glycosyltransferase WciJ (n=5), UDP-L-FucNAc Pathway FnlA (n=2), and capsule regulatory protein Wze (n=1) (Table 1 and Figure 3). These *cps* variants were detected in five GPSCs (GPSC26, 32, 55, and 334) and prevalence of 12F *cps* variants in these GPSCs were 0.6% (n=2/359), 2.4% (n=4/166), 2.0% (n=2/101) and 0.9% (n=1/101), respectively. It is of note that most disruptive mutations (6/9) arose from strand-slippage events supported by >100× sequence depth. For example, deletions of a single thymine in 5-bp homopolymer and adenine in 8-bp homopolymer disrupted *fnlA* in two GPSC32 isolates, and deletion of a single adenine in 8-bp homopolymer truncated *wciJ* in four different GPSC isolates.

**Table 1.** The serotype 12F capsule variants in *Streptococcus pneumoniae*.

| ERR | Disruptive mutation<br>(NT/AA)<sup>a</sup> | GPSC | Country | Year | Gene Function |
| --- | --- | --- | --- | --- | --- |
| SRR11152260 | *fnlA* 746delT<sup>b</sup><br>F250fs | 32 | USA | 2018 | UDP-L-FucNAc Pathway |
| SRR11152270 | *fnlA* 122delA<sup>c</sup><br>K43fs | 32 | USA | 2018 | UDP-L-FucNAc Pathway |
| ERR2091271 | *wciI* C107A<br>S36trunc | 55 | Israel | 2012 | Initial transferase |
| ERR2089444 | *wze* 640insAAT<br>K214trunc | 32 | Brazil | 2013 | Regulatory |
| ERR2090472 | *wciJ* 155insCTATCCTG<br>V75trunc | 26 | Nigeria | 2016 | Glycosyltransferase |
| ERR1065635 | *wciJ* 195delA<sup>c</sup><br>K54fs | 26 | Nepal | 2014 | Glycosyltransferase |
| ERR845755 | *wciJ* 195delA<sup>c</sup><br>K54fs | 32 | USA | 2007 | Glycosyltransferase |
| ERR2090766 | *wciJ* 195delA<sup>c</sup><br>K54fs | 55 | Israel | 2013 | Glycosyltransferase |
| DRR198456 | *wciJ* 195delA<sup>c</sup><br>K54fs | 334 | Japan | 2015-2017 | Glycosyltransferase |

<sup>a</sup>NT, mutations indicated in nucleotide sequence; AA, mutations indicated in amino acid sequence relative to the reference sequence of serotype 12F capsule encoding region (NCBI accession number CR931660). fs, Framshift; trunc, truncation
<sup>b</sup>Strand slippage mutations due to homopolymer of thymine (T) bases.
<sup>c</sup>Strand slippage mutations due to homopolymer of adenine (A) bases.

**Figure 3.**
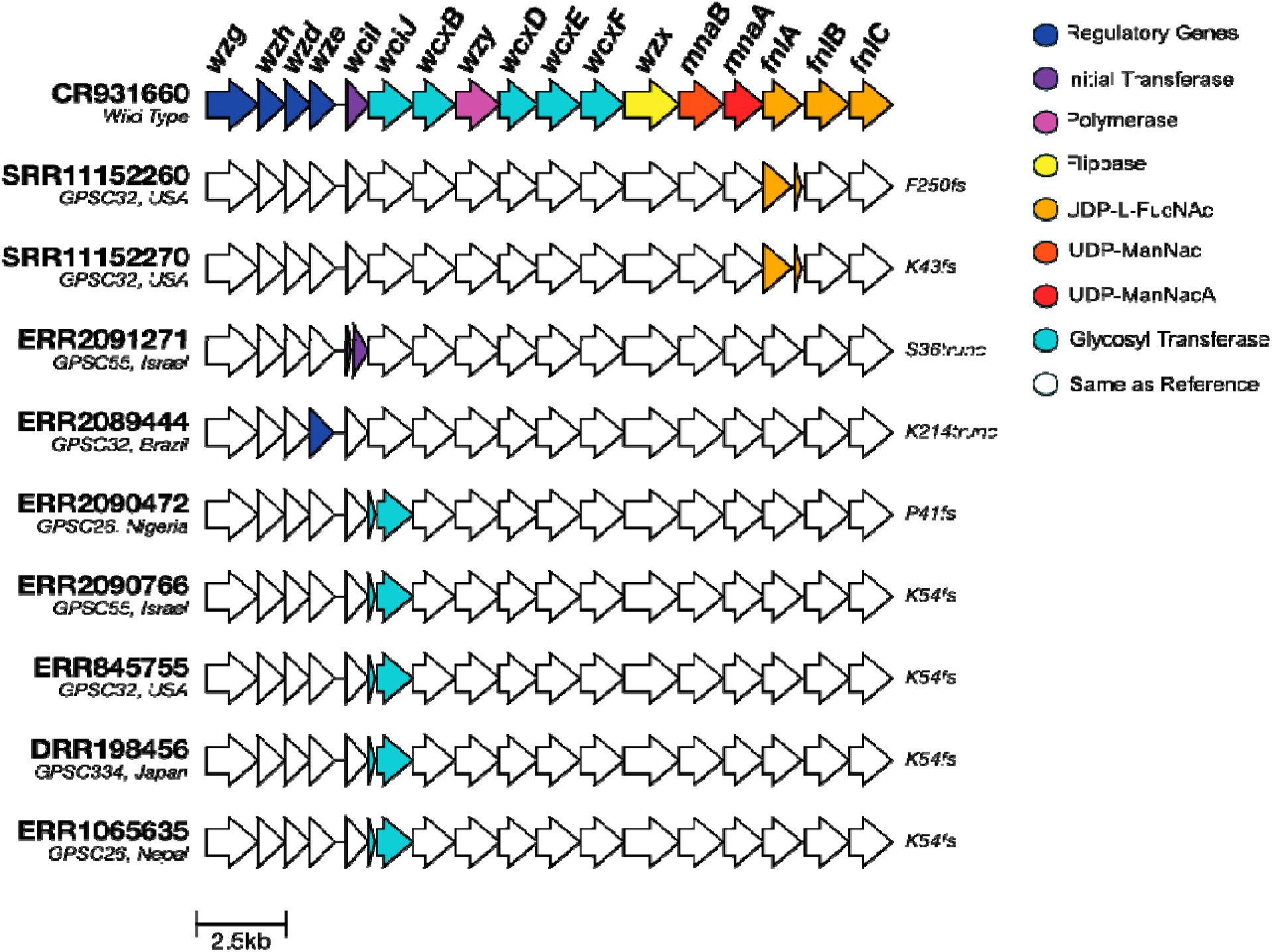
Genetic alignment of nine serotype 12F capsule variants.

Loss of WciL or WciJ, which are responsible for transferring the first and second sugars of the polysaccharide repeat unit, respectively, would likely prevent capsule formation.^52^ The strain in this study carries a truncated wze, resulting in the absence of a key tyrosine residue at the C-terminus; such deletions have previously been reported to lead to loss of capsule (Figure S7).^53^ This is due to the protein’s inability to autophosphorylate at the absent tyrosine residues. This consequentially impairs the interaction with the phosphatase Wzh, affecting the conformational cycle between phosphorylated and dephosphorylated states.^54^ In contrast, disruptive mutation observed in FnlA was also present in serotype 44, suggesting that it does not affect capsule production but likely the capsular structure.^55^ Further Quellung reaction using US CDC antisera confirmed the serotype 12F isolates with disruptive mutation in *fnlA* were serotype 44. WGS-based serotyping tools should be updated to more accurately recognise serotype 44.

Among the six *cps* variants, an identical disruptive mutation 195delA in *wciJ* was identified in four isolates from different lineages GPSC26, 32, 55 and 334 from Nigeria, USA, Israel and Japan respectively. The *wciJ* gene encodes a putative α-1,3-l-FucpNAc transferase to transfer α-L-FucpNAc following the initial sugar GalpNAc.^56^ The detected mutation occurred within an 8-bp adenine homopolymer, indicating a strand-slippage mutation. The deletion of an A into a 7-bp adenine homopolymer resulted in frameshift and pre-mature stop codon. However, this disruptive mutation did not abolish the capsule production as expected, because three of four isolates with Quellung isolates still typed as serotype 12F, indicating the presence of the capsule. Examining the read depth, we observed co-existence of the 8-bp adenine homopolymer (wildtype) and 7-bp adenine homopolymer (disrupted) in two strains, with wildtype-to-mutant ratios were 1:4 in the American isolate and 4:1 in the Japanese isolate based on the read depth. In contrast, the Israeli and Nepali isolates carried only the mutant form of *wciJ.* Apart from the Nepeali isolate which recovered from nasopharyngeal swab, the other three isolates were recovered from blood culture.

### Capsule production and vaccine protection of an evolutionarily convergent serotype 12F capsule variant

This heterogeneity prompted us to investigate whether the strain carrying the disruptive wciJ 195delA mutation is capable of capsule phase variation and its impact on vaccine-elicited protection. We constructed D39W strain with 12F *cps* (NUS0314), without *cps* (NUS0114), 12F *cps* with knockout *wciJ* (NUS7178), 12F *cps* with 195delA *wciJ* (NUS7324), and a backcross strain (NUS7389) derived from NUS7178 and NUS7324 (Table S2 and S3). Production of serotype 12F capsule was clearly observed in the D39W background, confirming that the D39W can express serotype 12F capsule (Figure 4). Deletion of *wciJ* abolished capsule formation as expected, whereas the *wciJ* (195delA) strains (NUS0314 and NUS7389) showed a mixed population of encapsulated and non-encapsulated pneumococci (Table S5). The presence of a capsule indicates that the encapsulated subpopulation possesses a functional *wciJ*. This is likely due to a reverting mutation from seven to eight adenines that restores WciJ functionality during replication. This observation is supported by both encapsulated and non-capsulated cells within a single chain, indicating rapid capsule phase variation (Figure S8-9). All the immunofluorescent assay images were collated in Figure S10-15.

**Figure 4.**
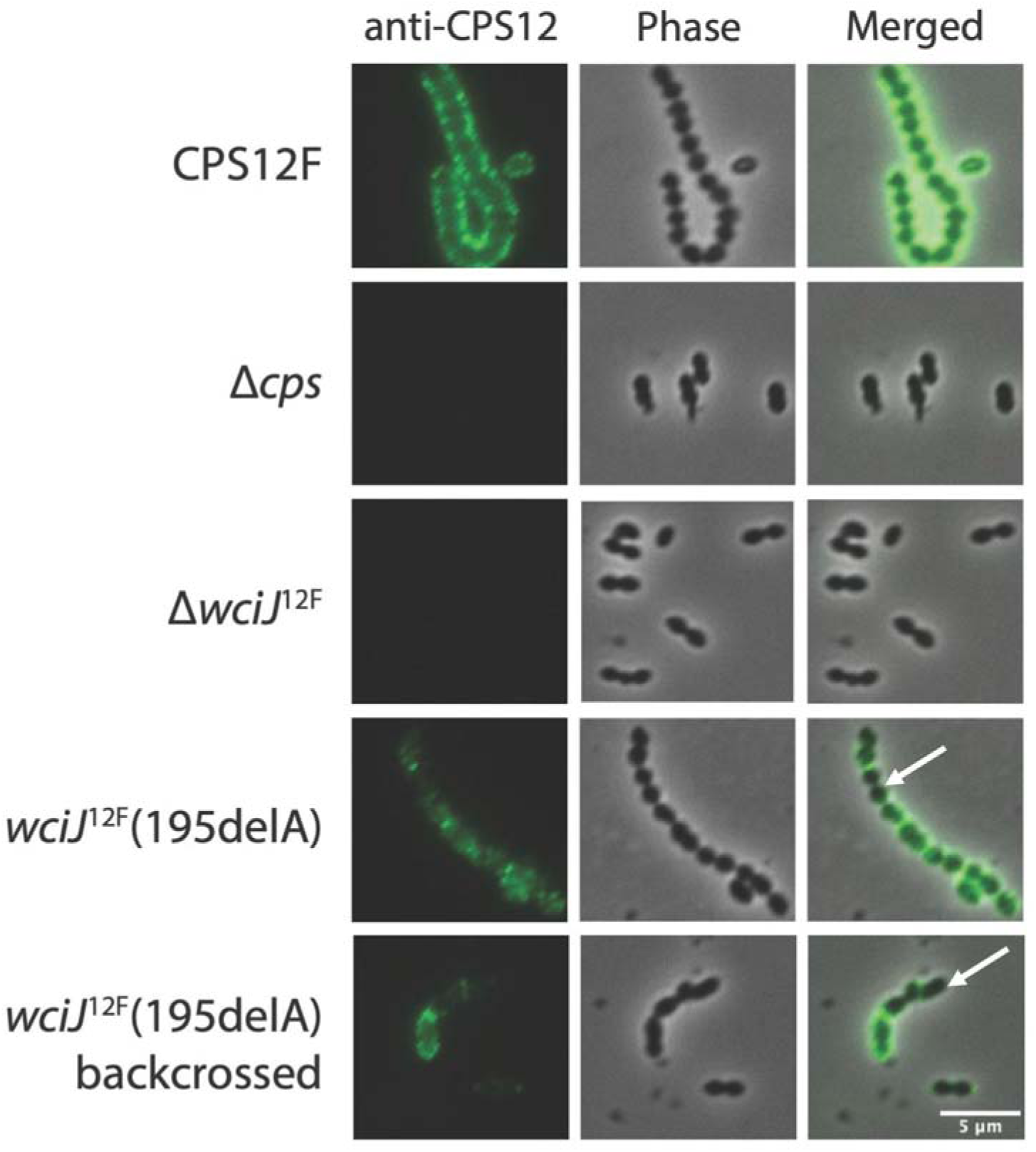
Immunofluorescence staining of serotype 12F capsule expressed by D39W strain. Strains were grown, centrifuged and stained with anti-serotype 12F CPS antisera. Anti-CPS antibodies were labelled with Alexa Fluor 488 secondary antibodies before imaging with UV fluorescence and phase contrast microscopy. Representative images from two biological replicates are shown. Scale bar, 5 µm. The white arrow shows the pneumococcal cells do not produce serotype 12F capsules.

To place this evolutionarily convergent event in a clinical context, we selected a pair of serotype 12F clinical strains with (GPS_NP_6940) and without (GPS_NP_2597) 195delA in *wciJ* otherwise highly similar genetically, differing by only 0.02% of nucleotides (391 SNPs across a 2,004,030 bp genome), to examine the vaccine-elicited protection using an opsonophagocytosis assay with sera collected from ten adults immunised with PPV23 (Figure 5). The OI of the 12F *cps* variant strain (geometric mean: 95.1, 95% CI: 17.3–524.1) was lower than that of the 12F *cps* wild-type strain (geometric mean: 252.8, 95% CI: 49.7–1286.3), although this difference was not statistically significant (p value = 0.63). A lower OI indicates that a higher concentration of antibodies is necessary to achieve 50% bacterial killing, suggesting that the 12F *cps* variant strain may exhibit increased resistance to serum-mediated killing. Notably, marked variation in OI was observed across both test strains and the reference strain (geometric mean: 360, 95% CI: 63.0-2064.0) likely reflecting heterogeneity among individual antisera.

**Figure 5.**
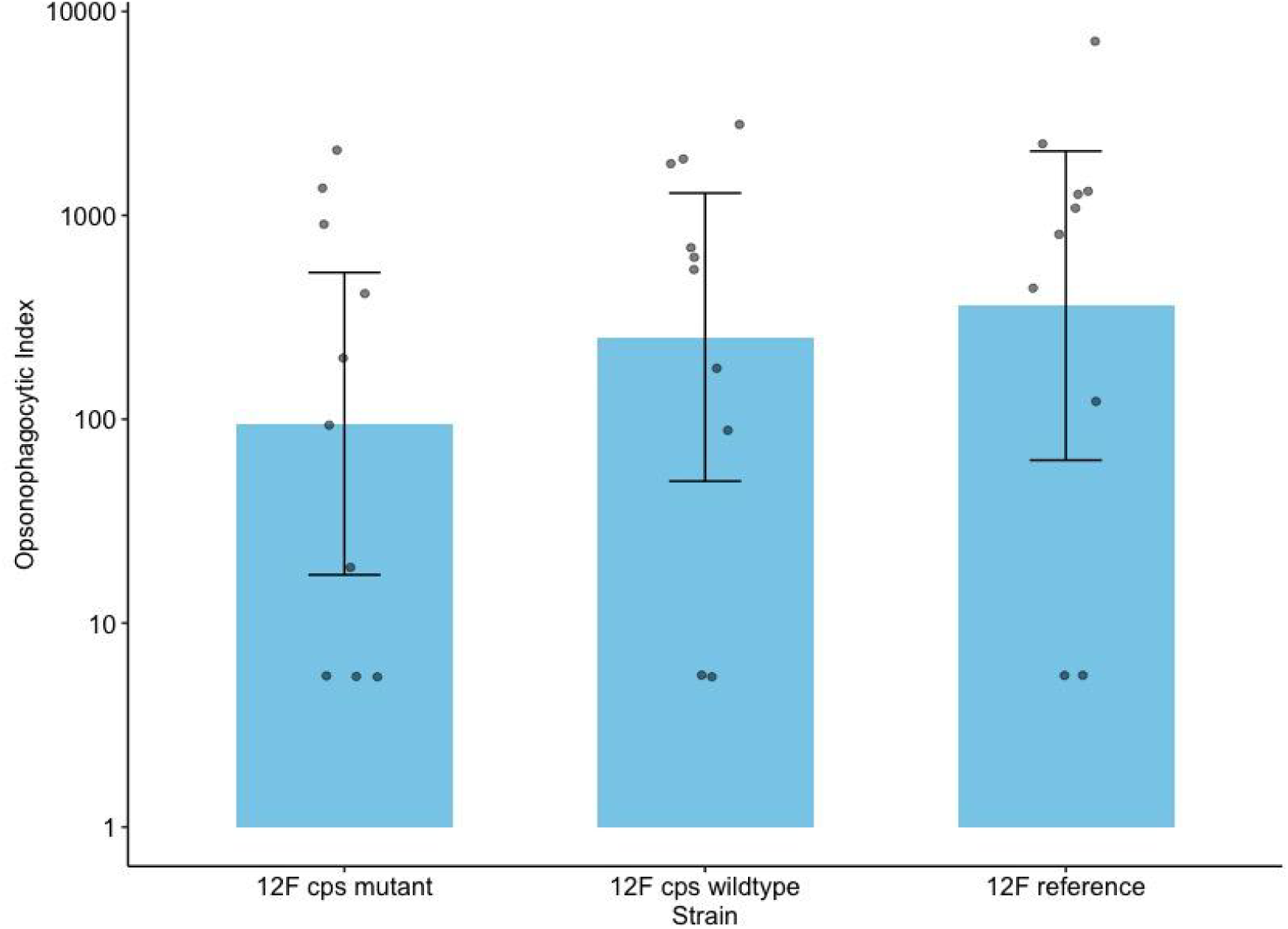
Opsonophagocytic index of the serotype 12F strains with 195delA in *wciJ* (GPS_NP_6940) and without the 195delA in *wciJ* (GPS_NP_2597), and 12F reference strain DS4009-06. A lower opsonophagocytic index indicates that a lower serum dilution is required to achieve 50% bacterial killing, suggesting increased resistance to antibody-mediated opsonophagocytic killing.

## Discussion

Our results indicated that the global increase in serotype 12F was driven by distinct pneumococcal lineages across different continents, and a globally-disseminated MDR lineage GPSC26. We identified previously unrecognised genetic diversity within the capsular encoding region of serotype 12F, providing evidence of convergent evolution in a glycosyltransferase WciJ. This convergent mutation in *wciJ* represents a novel mechanism for rapidly reversible on–off switching of capsule production through strand slippage mutation, leading to heterogeneity in capsule production in a single infection. This may reduce susceptibility to vaccine-elicited serum killing, as non-encapsulated subpopulations are inherently resistant to serotype-specific antibodies while remaining susceptible to innate immune clearance.^57^

Since the introduction of PCV13, GPSC26 has been increasingly identified, not only in Africa and Asia, but also more recently in the Netherlands,^13^ Spain,^58,59^ and Argentina (personal communication with Dr Nahuel Sanchez Eluchans), indicating its ongoing global spread. In addition to it being multidrug-resistant, GPSC26 also demonstrates a higher invasive disease potential, as compared with other serotype 12F lineages, such as GPSC32 and GPSC56.^60^ GPSC26 is also capable of transferring its 12F *cps* to other lineages and acquiring penicillin-resistance mutations through recombination. To date, no outbreaks caused by GPSC26 have been reported, whereas pan-susceptible lineages like GPSC32 have been linked to outbreaks among individuals experiencing homelessness in the US,^61^ and GPSC212 has been associated with outbreaks among shipyard workers in Finland.^29^

Unlike other continents, a variety of serotype 12F lineages were detected in Europe, including GPSC26, 32, 55, 212, and 334. The majority of serotype 12F isolates from Spain were GPSC26,^58,59^ while serotype 12F in the Netherlands is composed of GPSC26, GPSC32 and GPSC55.^13^ GPSC212 expressing serotype 12F has only been detected in Northern Europe including the Russia Federation^62^ and Finland.^29^ The recent licensure of PCV20 (Pfizer) in Europe is likely to change the landscape of serotype 12F IPD in Europe, and later in other continents. It is also of note that all the higher valency of PCVs include serotype 12F, including PCV21 V116, Merck), PCV24 (AFX3772, GSK; VAX-24, Vaxcyte), PCV25 (IVT-25, Inventprise), PCV31 (VAX-31, Vaxcyte).

This is the first study to map the genetic landscape of serotype 12F capsules, revealing the strand-slippage mutation as one of the major mechanisms driving capsule diversification that may result in variations in capsule production and structure. The same mechanism is also seen in serotypes 35B and 35D: when *wciG* is intact, the phenotype is serotype 35B, whereas disruption of *wciG* through strand slippage mutation shifts the phenotype to serotype 35D.^63^ The absence of the *wciG*-encoded acetyltransferase has been linked to high invasive disease potential.^63^ Similarly, strand slippage within a 7-bp adenine in *wciG* alters capsule phenotype from 33G to 33H.^64^ Beyond the *cps* region, strand slippage mutations in promoter region of *nanB* operon have also been observed, where they increase colonisation density and reduce mucus trapping.^65^ Future study with larger sample size could assess whether this deletion is directly associated with invasive disease progression. Notably, the detection of mixed wild-type and mutated *wciJ* alleles in serotype 12F isolates from blood cultures suggests that such mutations may represent ongoing microevolution within the host.

Capsule phase variation has previously been attributed to methylation^66^ and transcriptional factors influencing *cps* operon expression.^67,68^ Our findings revealed a novel mechanism mediated by strand slippage mutation in glycosyltransferase. Furthermore, this strand-slippage mutation in *wciJ* was not sporadic but occurred convergently across multiple lineages and countries over a ten-year period (2007–2017), strongly suggesting that it represents an adaptive mutation within the host. This was supported by previous experiments which showed that heterogeneity in capsule production plays an important role in the life cycle of pneumococci.^27^ While absence of the capsule enhances bacterial adherence, the presence of the capsule promotes survival under nutrient starvation and facilitates immune evasion.^27^ Collectively, this heterogeneous pneumococcal population is beneficial for colonisation and pathogenesis.^27^

Heterogeneity in capsule production within a pneumococcal population may also contribute to reduced susceptibility to opsonophagocytic killing, as non-encapsulated cells are inherently resistant to vaccine-elicited opsonophagocytosis. However, the non-encapsulated cells are still susceptible to innate immune clearance.^57^ It remains unclear whether this heterogeneous capsule production enhances population survival under vaccine-induced selective pressure.

The two serotype 12F isolates carrying frameshift and disruptive mutations in *fnlA* are actually serotype 44, as the serotype 44 reference *cps* sequence (NCBI accession number CR931717) also contains a disruptive mutation in *fnlA*. Although Mavroidi *et al*. previously reported that the only difference between serotypes 12F and 44 was a frameshift in *fnlC*,^56^ we believe this to be a typographical error. The current WGS-based serotyping methods such as PneumCaT, PneumoKITy and SeroBA v2.0 recognised nonsynonymous mutations at position L186P, L262I, G305A and A373T as the signatures of serotype 44. However, these nonsynonymous mutations were not observed in the Quellung typed serotype 44 isolates in this study but only seen in the serotype 44 *cps* reference sequence (NCBI accession number CR931717). It would be beneficial to update the WGS-based serotyping method to better recognise serotype 44 by detecting pre-mature stop codon in *fnlA*.

The lineage driving the increase in serotype 12F varies between countries, a pattern that may be partly explained by differences in antibiotic-selective pressures, similar to observations for another globally spreading lineage GPSC10 associated with the rise of serotype 24F.^34^ In Japan, where macrolide consumption is high (1,196 defined daily dose DDD per 1000,2023 estimation^69^) and penicillin use (<1 DDD per 1000) is low, the predominant 12F lineage is GPSC334, which is macrolide resistant but penicillinsusceptible (Table S3). In Israel, where penicillin use is high but macrolide use is relatively low, GPSC55 predominates and is penicillin resistant.^70^ GPSC26, which likely emerged in Africa and subsequently spread internationally, is resistant to cotrimoxazole—a drug widely used in Africa for HIV prophylaxis.^71^

This study has several limitations. The sampling strategy and collection timeframe varied across countries although the GPS project aimed to capture the changes before and after PCV introduction. For each strain tested, opsonophagocytic killing varied substantially between individual sera, resulting in reduced statistical power and a larger sample size of antisera would be required to detect the potential difference 12F variants. Despite these limitations, this comprehensive analysis provides an international perspective on serotype 12F lineages and capsules, highlighting a novel mechanism for capsule phase variation with potential impact on the PCV protection during the early stages of PCV20 implementation.

Our research further underscores the value of bacterial genome sequencing to elucidate pneumococcal lineage dynamics during PCV implementation and, when integrated with experimental validation, to uncover mechanisms of capsule production heterogeneity and their potential contribution to vaccine evasion.

## The Global Pneumococcal Sequencing Consortium authors

Patrick E Akpaka, Maaike Alaerts, Mushal Ali, Samanta Cristine Grassi Almeida, Martin Antonio, Houria Belabbès, Rachel Benisty, Godfrey Bigogo, Abdullah W Brooks, Philip E Carter, Kulkanya Chokephaibulkit, Piriyaporn Chongtrakool, Stuart C Clarke, Jennifer Cornick, Alejandra Corso, Nicholas Croucher, Ron Dagan, Alexander Davydov, Idrissa Diawara, Sanjay Doiphode, Ekaterina Egorova, Naima Elmdaghri, Nahuel Sanchez Eluchans, Özgen Köseoglu Eser, Dean B Everett, Diego Faccone, Ebenezer Foster-Nyarko, Paula Gagetti, Noga Givon-Lavi, Anne von Gottberg, Linda De Gouveia, Md Hasanuzzaman, Pak Leung Ho, Waleria Hryniewicz, Margaret Ip, Rama Kandasamy, Tamara Kastrin, Jeremy Keenan, Anmol Kiran, Keith P Klugman, Brenda Kwambana-Adams, Claudia Lara, Pierra Law, Deborah Lehmann, Cebile Lekhuleni, Shabir A. Madhi, Benild Moiane, Geetha Nagaraj, Kedibone Ndlangisa, Michele Nurse-Lucas, Susan A Nzenze, Stephen Obaro, Theresa J. Ochoa, Wanatpreeya Phongsamart, Mignon du Plessis, Nurit Porat, Weronika Puzia, KL Ravikumar, Gail Rodgers, Ewa Sadowy, Samir K. Saha, Senjuti Saha, Eric Sampane-Donkor, Shamala Devi Sekaran (Sekaran SD), Betuel Sigauque, Anna Skoczynska, Somporn Srifuengfung, Hoa Ngo Thi, Peggy-Estelle Tientcheu, Leonid Titov, Paul Turner, Yulia Urban, Balaji Veeraraghavan, Elena Voropaeva, Hui Wang, Nicole Wolter, Khalid Zerouali, Chunjiang Zhao, and Jonathan Zintgraff.

## Acknowledgement

The study was cofunded by the Gates Foundation (grant code INV-003570) and the Wellcome Sanger Institute (core Wellcome grants 098051 and 206194). We extend our sincere gratitude to all members of the Global Pneumococcal Sequencing Consortium for their contributions of sample collection, processing, and collaborative efforts. We also appreciate the valuable feedback from Prof Mark Holmes at University of Cambridge and Dr David Litt in UKHSA who had reviewed this study as part of Thi N M Huynh’s MPhil degree at the University of Cambridge. We appreciate Dr Raquel Sá-Leão’s comments on this work when Dr Stephanie Lo presented it at ITQB NOVA. Additionally, we acknowledge the support from the sequencing facility and the Pathogen Informatics team at the Wellcome Sanger Institute, in Hinxton, UK. The findings and conclusions detailed in this manuscript are those of the authors and do not necessarily represent the official position of the US Centers for Disease Control and Prevention.

