## Supplementary material for "Global population structure and phase variation of serotype 12F *Streptococcus pneumoniae* following the introduction of pneumococcal conjugate vaccine": Figure S1

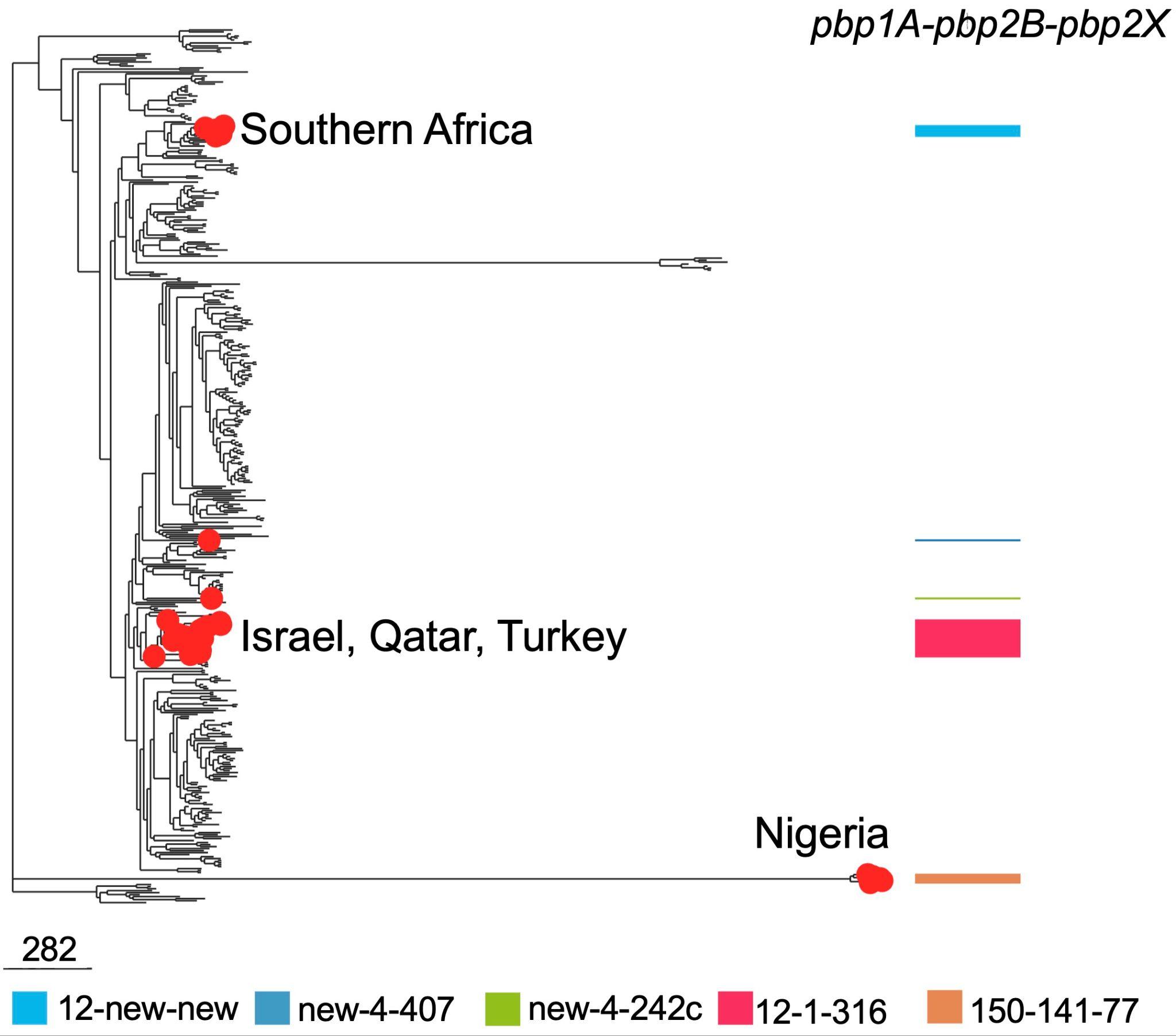

**Figure S1 Global phylogeny of GPSC26 with penicillin-resistant isolates highlighted and overlaid with penicillin-binding protein (*pbp*) profile. The scale bar refers to the number of single nucleotide polymorphisms.**
