## Supplementary material for "Global population structure and phase variation of serotype 12F *Streptococcus pneumoniae* following the introduction of pneumococcal conjugate vaccine": Figure S2

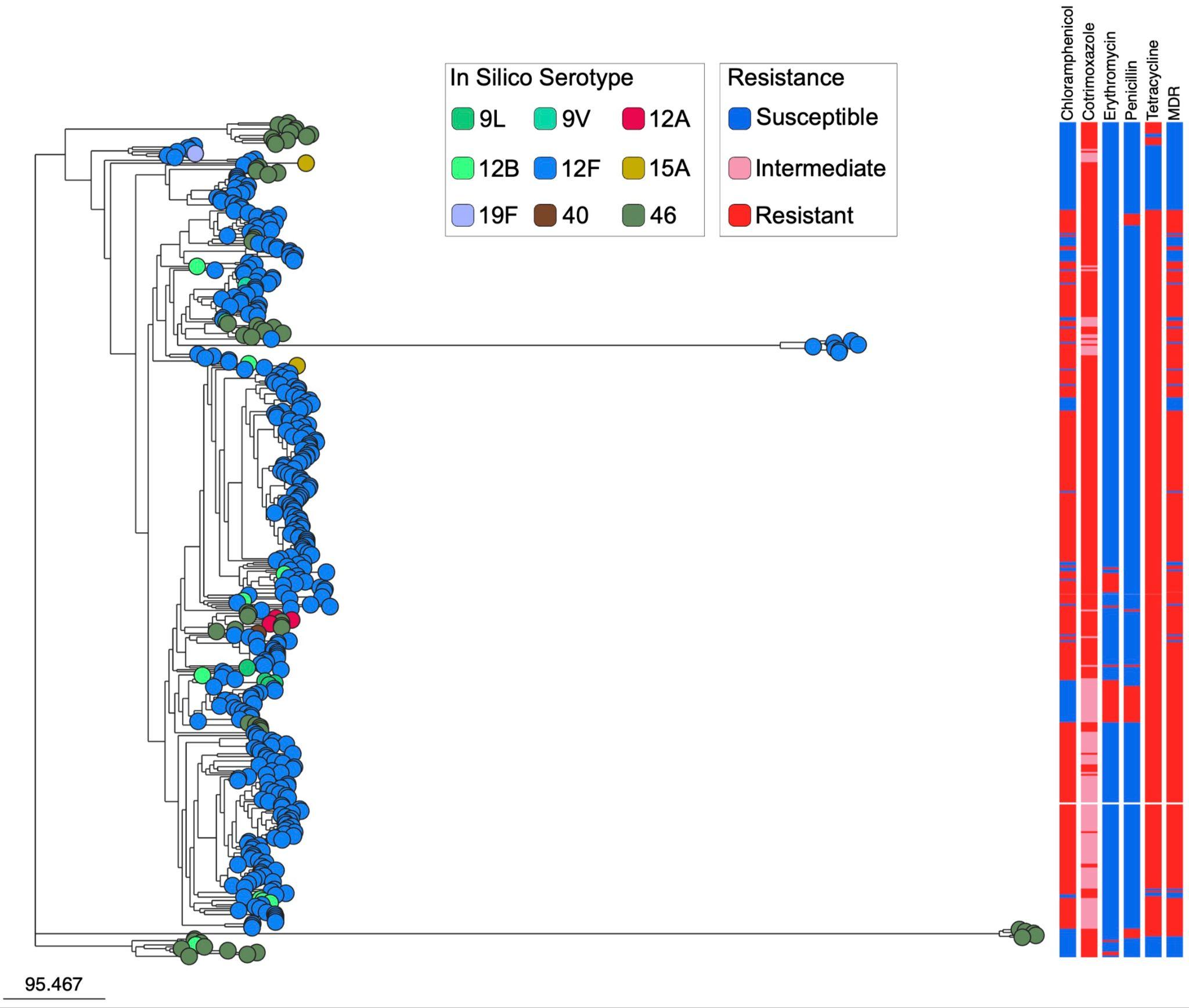

**Figure S2 Global phylogeny of GPSC26 with nodes coloured by serotypes, overlaid with antimicrobial resistance profile. The scale bar refers to the number of single nucleotide polymorphisms.**
