## Supplementary material for "Global population structure and phase variation of serotype 12F *Streptococcus pneumoniae* following the introduction of pneumococcal conjugate vaccine": Figure S3

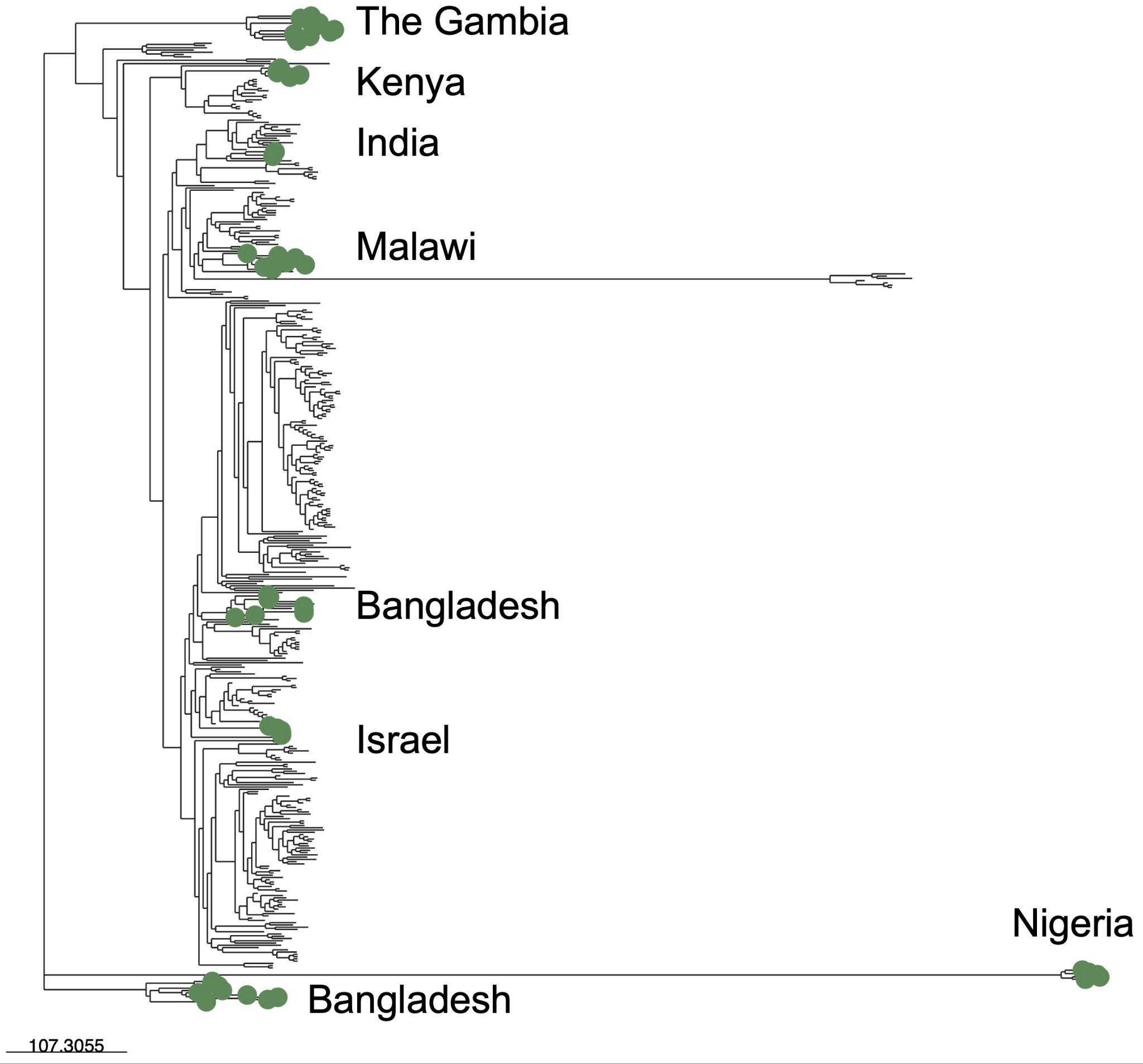

**Figure S3 Global phylogeny of GPSC26 with serotype 46 isolates highlighted. The scale bar refers to the number of single nucleotide polymorphisms.**
