## Supplementary material for "Global population structure and phase variation of serotype 12F *Streptococcus pneumoniae* following the introduction of pneumococcal conjugate vaccine": Figure S4

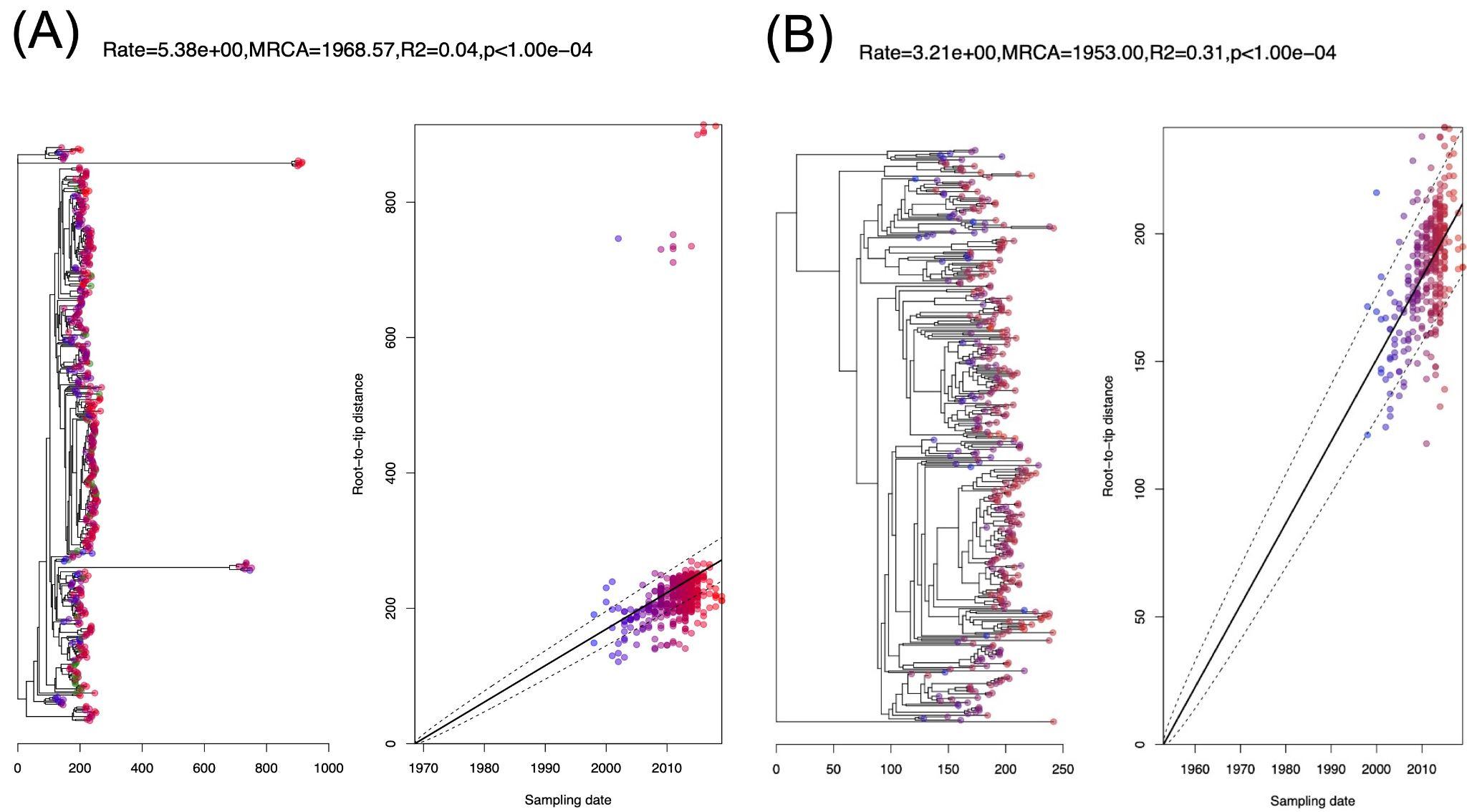

**Figure S4 Phylogenies and linear regression of root-to-tip distance against year of collection with (A) and without (B) outliners.** The exclusion of outliers improved the temporal signal, increasing the R^2^ value from 0.04 to 0.31, which enabled convergence in the BEAST analysis, indicated by the effective sample size (ESS) >200.
