## Supplementary material for "Global population structure and phase variation of serotype 12F *Streptococcus pneumoniae* following the introduction of pneumococcal conjugate vaccine": Figure S5

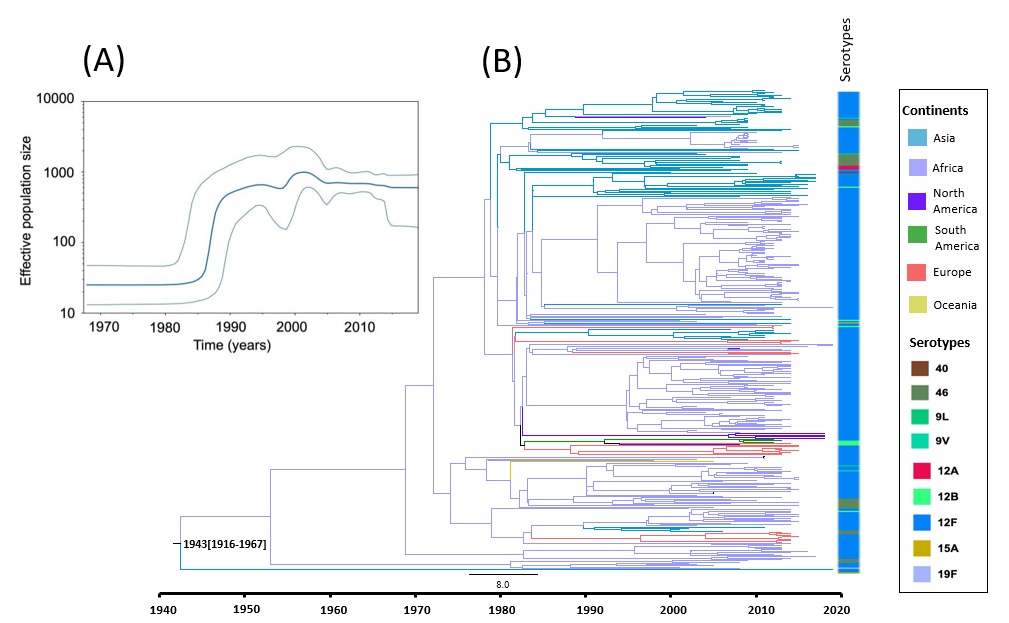

**Figure S5 Globally-spreading serotype 12F pneumococcal lineage GPSC26 (n=398).** (A) Bayesian skyline plot of estimated median effective population size of Streptococcus pneumoniae GPSC26 over time; (B) A time-resolved GPSC26 phylogeny coloured by continents; Asia (blue); Africa (light purple); North America (dark purple); South America (green); Europe (orange); Oceania (yellow).
