## Supplementary material for "Global population structure and phase variation of serotype 12F *Streptococcus pneumoniae* following the introduction of pneumococcal conjugate vaccine": Figure S6

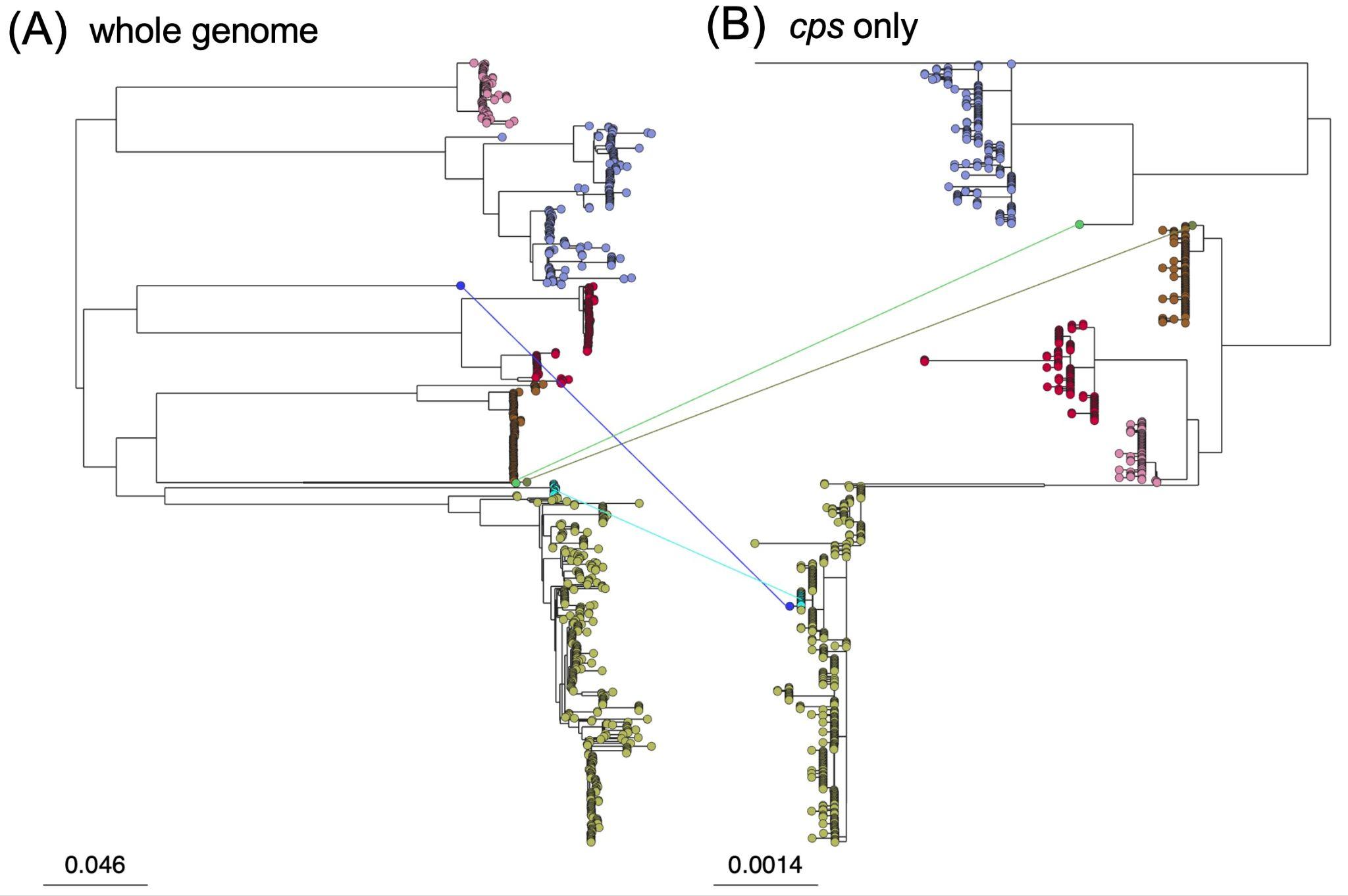

**Figure S6 Phylogenies of serotype 12F isolates built upon single nucleotide polymorphism across the genome (A) and capsular encoding region only (B).** The lines between the phylogenies indicated the four potential capsular switching events. For example, GPSC26 (olive green) are the potential donor of *cps* for GPSC648 (blue) and GPSC212 (turquoise) as their cps genetic sequences are highly similar though the genetic background were distantly related. The scale bar refers to a genetic distance of nucleotide substitutions per site.
