## Supplementary material for "Global population structure and phase variation of serotype 12F *Streptococcus pneumoniae* following the introduction of pneumococcal conjugate vaccine": Figure S7

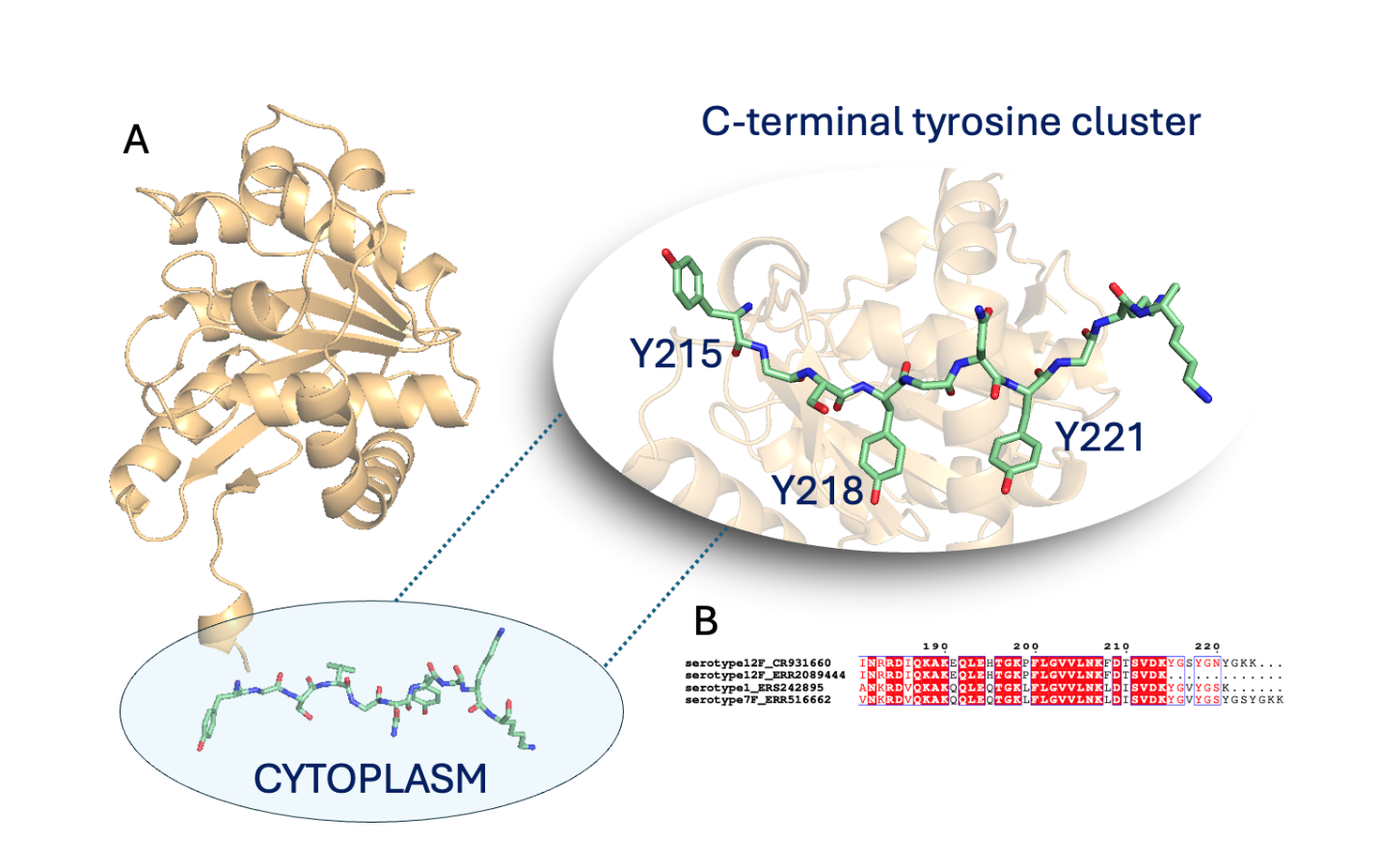

**Figure S7** (A) Predicted protein structure of 12F Wze by AlphaFold (CR931660). The residues essential for the phosphorylation reaction are shown in green sticks, coloured by atom type. (B) Aligned sequences from serotype 12F, 12F (Variant ERR2089444), serotype1, serotype 7F, respectively. The truncated sequence of Variant ERR2089444 in this study is missing the key tyrosine residues.
