## Supplementary material for "Global population structure and phase variation of serotype 12F *Streptococcus pneumoniae* following the introduction of pneumococcal conjugate vaccine": Figure S8

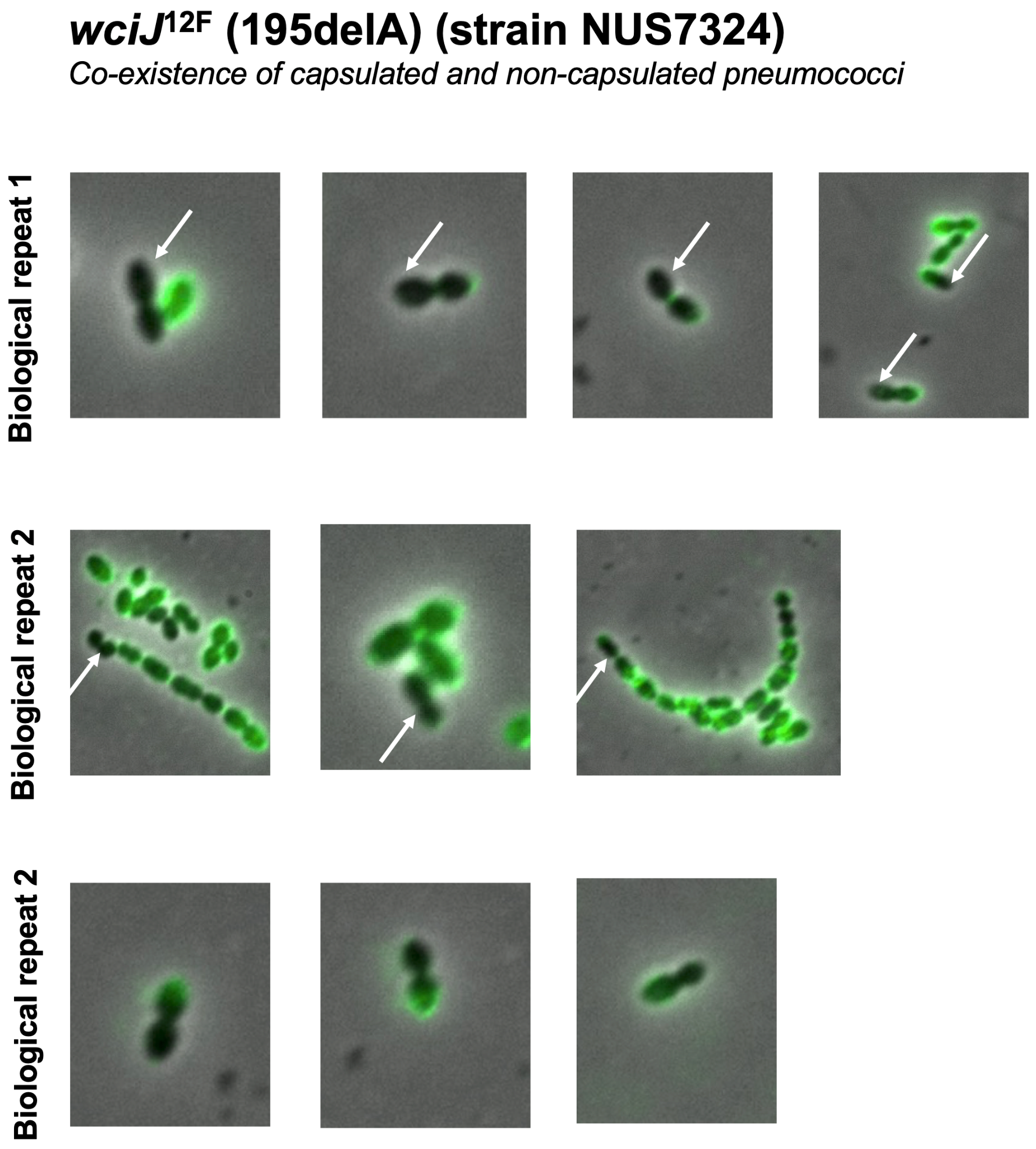

**Figure S8 Coexistence of encapsulated and non-encapsulated pneumococcal cells in a single chain observed from the D39W strain expressing the serotype 12F capsule encoding region with 195delA mutation in *wciJ* gene (NUS7324).**
