## Supplementary material for "Global population structure and phase variation of serotype 12F *Streptococcus pneumoniae* following the introduction of pneumococcal conjugate vaccine": Figure S9

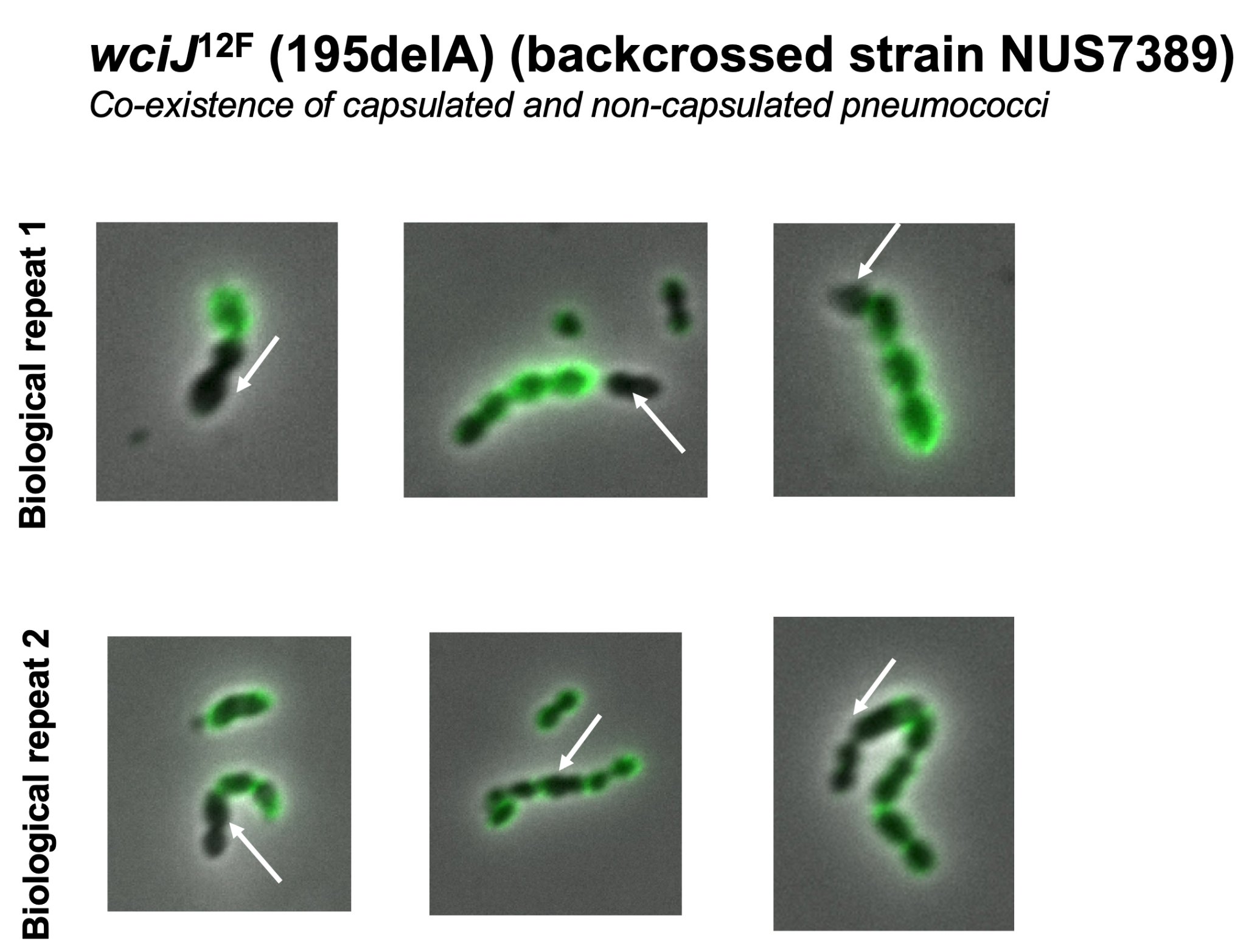

**Figure S9 Coexistence of encapsulated and non-encapsulated pneumococcal cells in a single chain observed from the backcross strain (NUS7389) derived from NUS7178 (D39W-12F*cps*-∆*wciJ*) and NUS7324 (D39W-12F*cps*-195del-*wciJ*).**
