## Supplementary material for "Global population structure and phase variation of serotype 12F *Streptococcus pneumoniae* following the introduction of pneumococcal conjugate vaccine": Figure S10

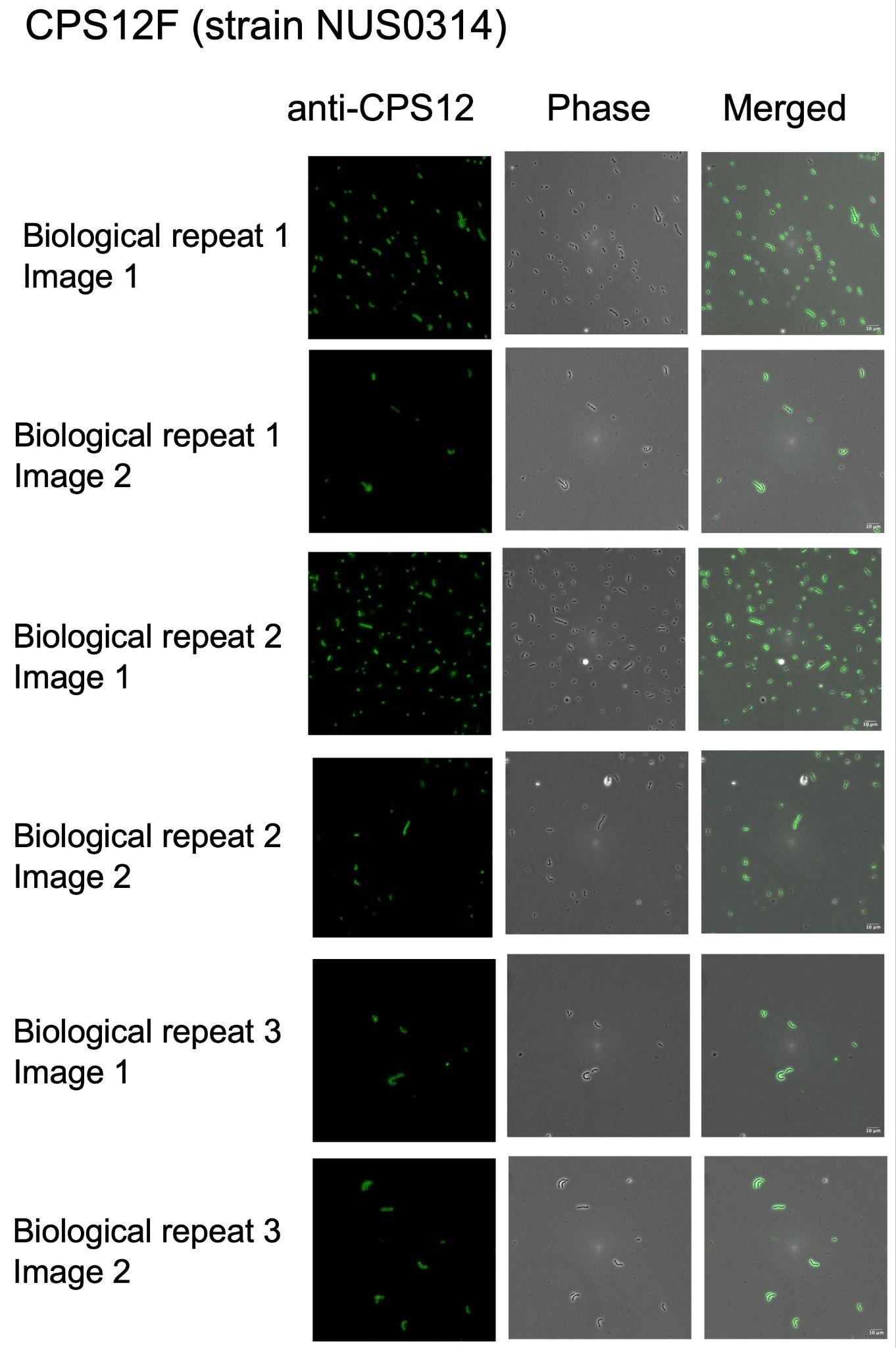

**Figure S10 Immunofluorescence staining of the D39W strain expressing the serotype 12F capsule (NUS0314) from clinical isolate PATH122 (US CDC).**
