## Supplementary figures and images for "Global population structure and phase variation of serotype 12F *Streptococcus pneumoniae* following the introduction of pneumococcal conjugate vaccine"

### Figure S11

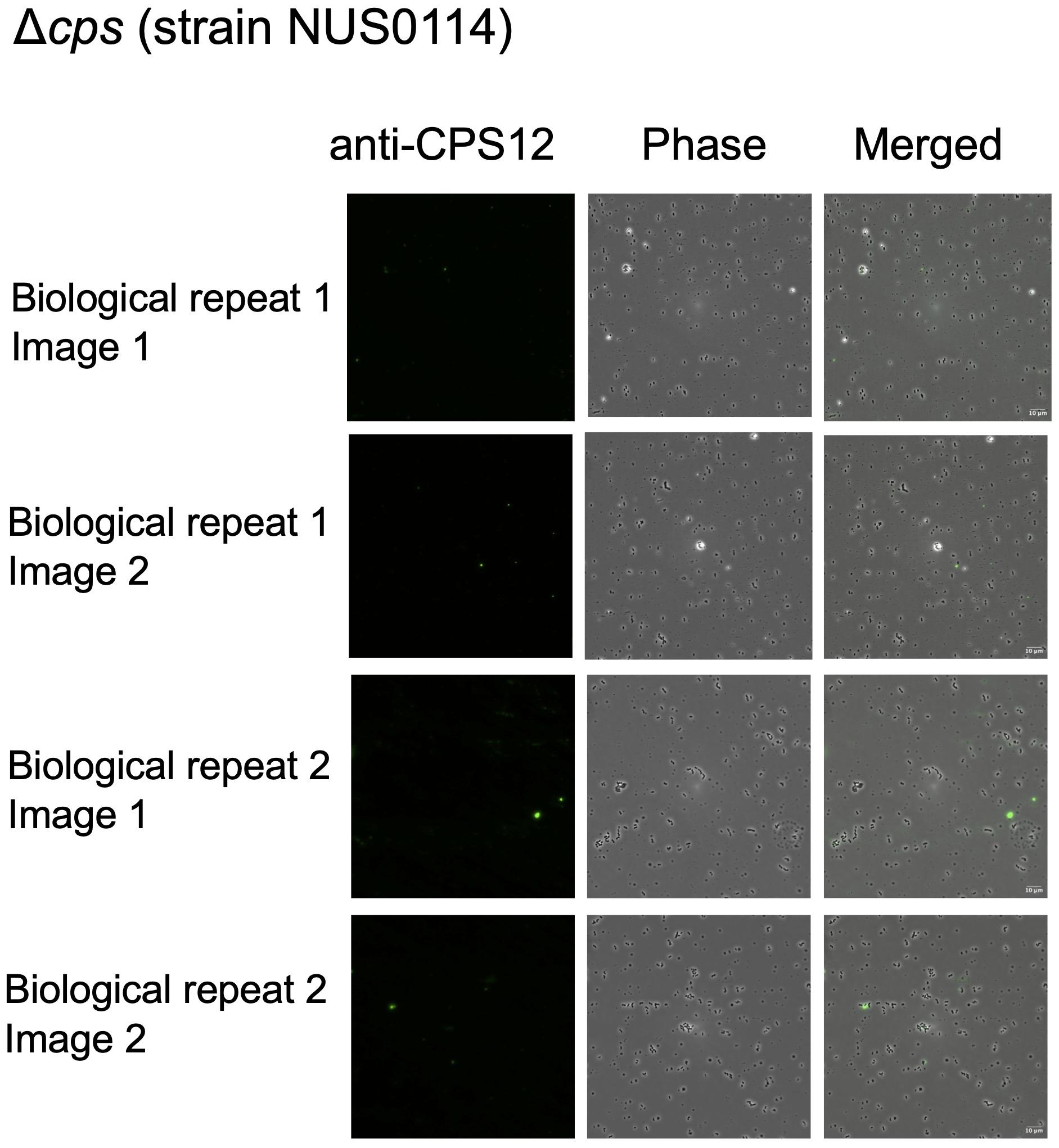

**Figure S11 Immunofluorescence staining of the D39W strain without capsule encoding region (NUS0114).**
