## Supplementary material for "Global population structure and phase variation of serotype 12F *Streptococcus pneumoniae* following the introduction of pneumococcal conjugate vaccine": Figure S12

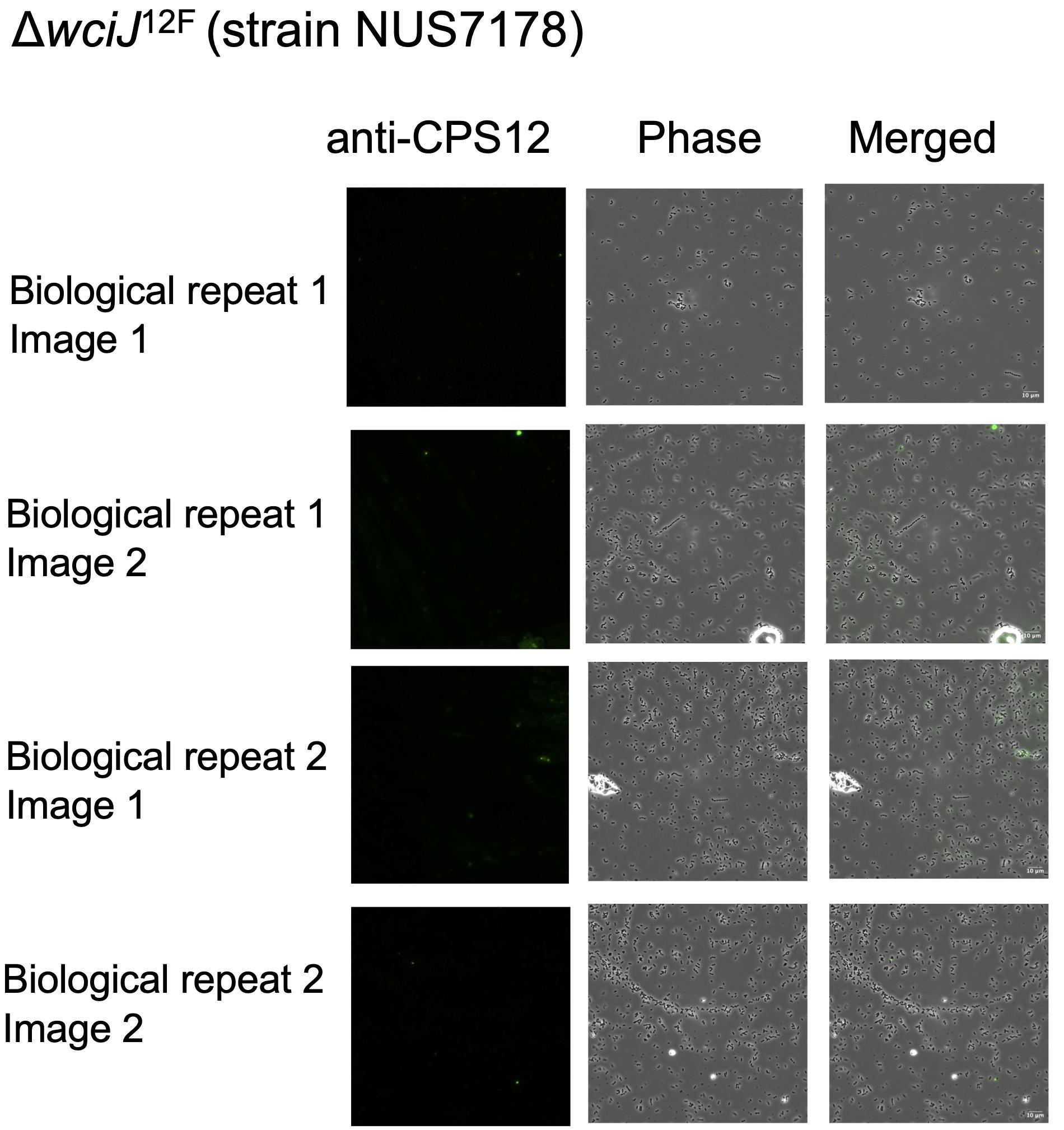

**Figure S12 Immunofluorescence staining of the D39W strain expressing the serotype 12F capsule encoding region with the wciJ gene knocked out (NUS7178).**
