## Supplementary material for "Global population structure and phase variation of serotype 12F *Streptococcus pneumoniae* following the introduction of pneumococcal conjugate vaccine": Figure S14

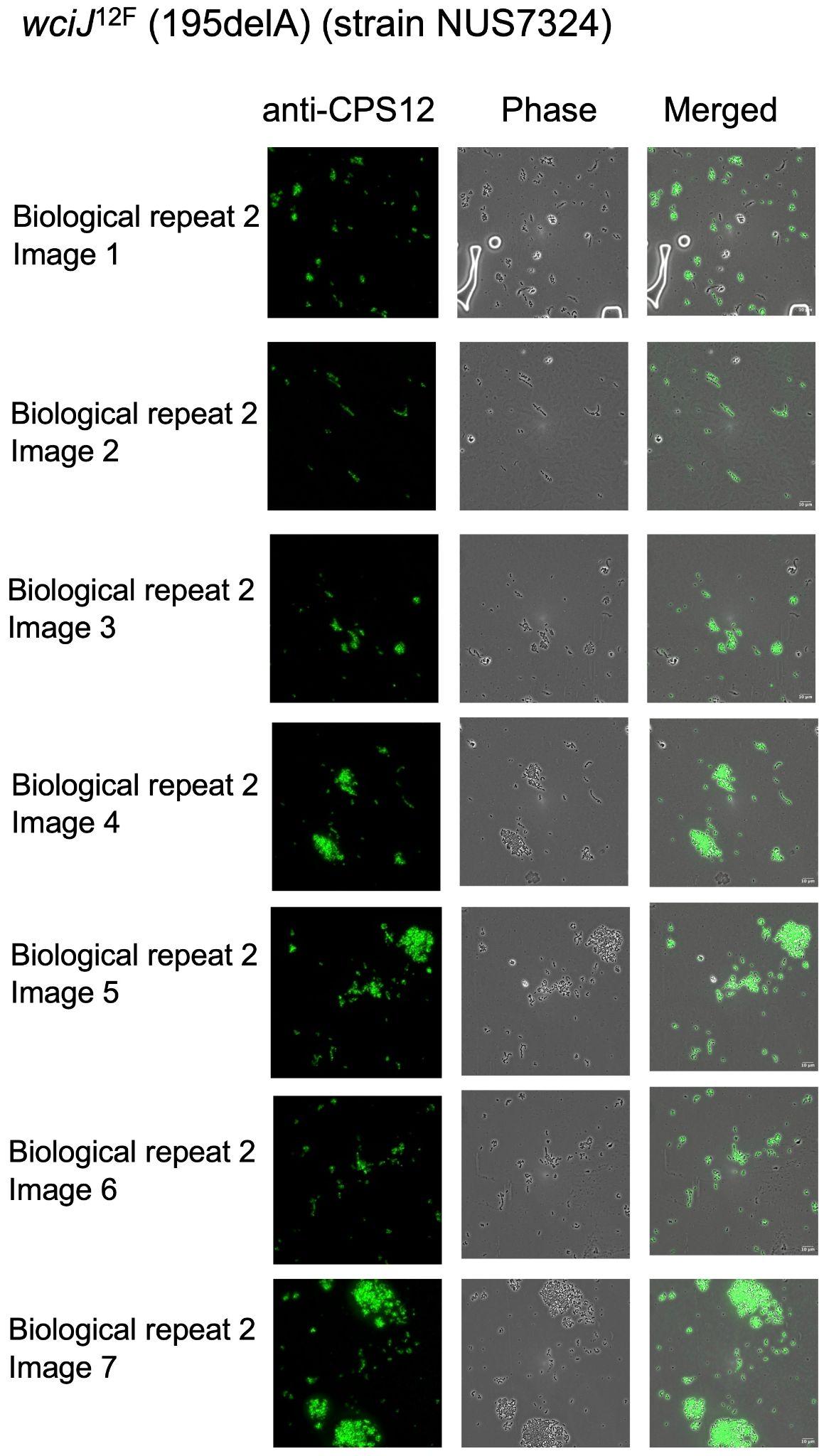

**Figure S14 Immunofluorescence staining of the D39W strain expressing the serotype 12F capsule encoding region with 195delA mutation in *wciJ* gene (NUS7324). The figure summarised six images from the 2nd biological replicate.**
