## Supplementary material for "Global population structure and phase variation of serotype 12F *Streptococcus pneumoniae* following the introduction of pneumococcal conjugate vaccine": Table S1

Table S1. Serotype 12F pneumococcal collections from literature review and Global Pneumococcal Sequencing (GPS) project database

| **No.** | **Sources of collection** | **No. of isolates** | **Country of origin** | **PCV**  **introduction** | **Years of study** | **References** |
| --- | --- | --- | --- | --- | --- | --- |
| 1 | Outbreak of invasive pneumococcal disease among shipyard workers, Turku, Finland, May to November 2019 | 9 | Finland | 2010(PCV10) | 2019 | (Linkevicius et al., 2019) |
| 2 | Local outbreak of Streptococcus pneumoniae serotype 12F caused high morbidity and mortality among children and adults | 6 | Japan | 2010(PCV7) | 2016 | (Ikuse et al., 2018) |
| 3 | Assessment of the local clonal spread of Streptococcus pneumoniae serotype 12F caused invasive pneumococcal diseases among children and adults | 14 | Japan | 2010(PCV7) | 2016-2018 | (Nakanishi et al., 2019) |
| 4 | Streptococcus pneumoniae Serotype 12F-CC4846 and Invasive Pneumococcal Disease after Introduction of 13-Valent Pneumococcal Conjugate Vaccine, Japan, 2015-2017 | 74 | Japan | 2010(PCV7) | 2015-2017 | (Nakano et al., 2020a) |
| 5 | Invasive Pneumococcal Strain Distributions and Isolate Clusters Associated With Persons Experiencing Homelessness During 2018 | 79 | United States | 2000(PCV7) | 2018 | (Metcalf et al., 2021) |
| 6 | Global Pneumococcal Sequence Project Database (last accessed on 25^th^ March 2022) | 11 | Argentina | 2012(PCV13) | 1998-1999,  2010-2013 | (Gagetti et al., 2021) |
| 7 |  | 12 | Bangladesh | 2015(PCV10) | 2004-2017 | GPS |
| 8 |  | 1 | Belarus | 2014(PCV10) | 2014 | GPS |
| 9 |  | 16 | Brazil | 2010(PCV10) | 2008-2009,  2012-2013 | (Almeida et al., 2021) |
| 10 |  | 1 | Cambodia | 2015(PCV13) | 2013 | GPS |
| 11 |  | 1 | Cameroon | 2011(PCV13) | 2013 | GPS |
| 12 |  | 1 | Ecuador | 2010(PCV7) | 2002 | GPS |
| 13 |  | 1 | Ethiopia | 2011(PCV10) | Unknown | GPS |
| 14 |  | 4 | Ghana | 2012(PCV13) | 2011, 2019 | GPS |
| 15 |  | 1 | Guatemala | 2012(PCV13) | 2002 | GPS |
| 16 |  | 1 | Hungary | 2009(PCV7) | 2005 | GPS |
| 17 |  | 12 | India | 2017(PCV13) | 2001-2017 | (Nagaraj et al., 2021) |
| 18 |  | 101 | Israel | 2009(PCV13) | 2006-2014 | (Lo et al., 2019) |
| 19 |  | 21 | Kenya | 2011(PCV10) | 2003, 2012-2016 | GPS |
| 20 |  | 35 | Malawi | 2011(PCV13) | 2003-2015 | (Lo et al., 2019) |
| 21 |  | 3 | Mongolia | 2016(PCV13) | 2003-2004, 2006 | GPS |
| 22 |  | 3 | Mozambique | 2013(PCV10) | 2008-2009 | GPS |
| 23 |  | 4 | Nepal | 2015(PCV10) | 2012, 2014, 2017 | GPS |
| 24 |  | 30 | Netherlands | 2006(PCV10) | 2012-2015 | GPS |
| 25 |  | 1 | Niger | 2014(PCV13) | Unknown | GPS |
| 26 |  | 2 | Nigeria | 2014(PCV10) | 2016 | GPS |
| 27 |  | 1 | Pakistan | 2012(PCV10) | 2013 | GPS |
| 28 |  | 1 | Papua New Guinea | 2013(PCV13) | 1990 | (Mellor et al., 2022) |
| 29 |  | 4 | Peru | 2009(PCV7) | 2008, 2010-2011 | (Hawkins et al., 2017) |
| 30 |  | 3 | Poland | 2017(PCV7) | 2008-2010 | GPS |
| 31 |  | 21 | Qatar | 2005(PCV7)  2010(PCV13) | 2010-2014 | GPS |
| 32 |  | 3 | Russian Federation | 2014(PCV13) | 2016-2017 | (Egorova et al., 2022) |
| 33 |  | 2 | Senegal | 2013(PCV13) | 2011-2012 | GPS |
| 34 |  | 140 | South Africa | 2009(PCV7)  2011(PCV13) | 2000-2014 | (Lo et al., 2019) |
| 35 |  | 1 | Thailand | 2013(PCV13) | 2011 | GPS |
| 36 |  | 100 | The Gambia | 2009(PCV7)  2011(PCV13) | 2002-2016 | (Lo et al., 2019) |
| 37 |  | 9 | Togo | 2014(PCV13) | 2011-2012, 2014 | GPS |
| 38 |  | 1 | Turkey | 2008(PCV7) | 2006 | GPS |
| 39 |  | 1 | Unknown | N/A | Unknown | GPS |
| 40 |  | 65 | United States | 2000(PCV7)  2010(PCV13) | 1998-2015 | GPS |
| 41 |  | 10 | West Africa |  | Unknown | GPS |
| **Total** |  | 806 |  |  |  |  |
