## Supplementary material for "Global population structure and phase variation of serotype 12F *Streptococcus pneumoniae* following the introduction of pneumococcal conjugate vaccine": Table S2

Table S2 Strains used to confirm the serotype 12F capsule production

| **Strain** | **Relevant genotype^a,b^** | **Derivation** | **Selectable marker^c^** | **Source** |
| --- | --- | --- | --- | --- |
| IU1690 | Serotype 2 strain D39W | - | - | (Lanie et al., 2007) |
| IU1781 | *rpsL1* | *rpsL1* x IU1690 | Str^R^ | (Kazmierczak et al., 2009) |
| PATH122 | Serotype 12F clinical isolate | - | - | CDC, USA |
| NUS0114 | *rpsL1 ∆cps*::P-*sacB*-*kan*-*rpsL*^+^ | *∆cps*::P-*sacB*-*kan*-*rpsL^+^* x IU1781 | Kan^R^, Str^S^, Suc^S^ | (Chua et al., 2021) |
| NUS0195 | *rpsL1* CPS12F | *cps* of PATH122 x NUS0114 | Str^R^ | (Chun et al., 2023) |
| NUS0314 | *rpsL1* CPS12F  (backcrossed) | *cps* of NUS0195 x NUS0114 | Str^R^ | (Chun et al., 2023) |
| NUS7178 | *rpsL1* CPS12F ∆*wciJ*::P-*spec*-*rpsL^+^* | *∆wciJ*::P-*spec*-*rpsL^+^* x NUS0314 | Spec^R^, Str^S^ | This study |
| NUS7324 | *rpsL1* CPS12F *wciJ*(195delA) | *wciJ*(195delA) x NUS7178 | Str^R^ | This study |
| NUS7389 | *rpsL1* CPS12F *wciJ*(195delA)  (backcrossed) | *wciJ*(195delA) of NUS7324 x NUS7178 | Str^R^ | This study |

^a^Strains were constructed as described in Methods. “::” indicates an insertion into a gene. Unless otherwise specified, deletion mutants retain 30 base pairs 5’ and 3’ of the corresponding open reading frames to avoid polarity. To select for the strains, antibiotics were used at final concentrations of 150 µg/ml for spectinomycin (Spec) or 300 µg/ml for streptomycin (Str) (Sigma-Aldrich).

^b^Primers used to synthesize fusion amplicons are listed in Supplementary Table S2.

^c^Antibiotic resistance markers: StrR, streptomycin; KanR, kanamycin; SucS, sucrose sensitive (sacB+), SpecR, spectinomycin resistant.
