## Supplementary material for "Global population structure and phase variation of serotype 12F *Streptococcus pneumoniae* following the introduction of pneumococcal conjugate vaccine": Table S3

Table S3 Primers used to synthesize fusion amplicons to construct the strains in table S2

| **Primer** | **Sequence (5’ to 3’)** | **Template** | **Amplicon** |
| --- | --- | --- | --- |
| **For construction of ∆*wciJ::*P-*sacB*-*kan*-*rpsL*^+^** | | | |
| N2658 | GAATTGTAATCATTAGTGTTCATACCAAG | PATH122 | *wciJ* 5’ |
| N2659 | CATTATCCATTAAAAATCAAACGGATCCTAACAAACAAATAGAATTTTCATTGCATC |  |  |
| N1 | TAGGATCCGTTTGATTTTTAATGGATAATG | NUS0092 | P-*spec*-*rpsL^+^* |
| N2 | GGGCCCCTTTCCTTATGCTTTTG |  |  |
| N2660 | CAAAAGCATAAGGAAAGGGGCCCTCGAAAGAGTGGTTTATGACATACTTAGAAAATC | PATH122 | *wciJ* 3’ |
| N2661 | GAGCGGCTATTAGCCTCGGTTG |  |  |
| For construction of *wciJ*(156delA) | | | |
| N2658 | GAATTGTAATCATTAGTGTTCATACCAAG | PATH122 | *wciJ*(195delA) 5’ |
| N3153 | GTCTCTCTTCTTTTTTTCTCTTAC |  |  |
| N3152 | GTAAGAGAAAAAAAGAAGAGAGAC | PATH122 | *wciJ*(195delA) 3’ |
| N2661 | GAGCGGCTATTAGCCTCGGTTG |  |  |
| For construction of *wciJ*(156delA) full fragment | | | |
| N2658 | GAATTGTAATCATTAGTGTTCATACCAAG | NUS7324 | *wciJ*(195delA) |
