## Supplementary material for "Global population structure and phase variation of serotype 12F *Streptococcus pneumoniae* following the introduction of pneumococcal conjugate vaccine": Table S4

Table S4 Common antimicrobial resistance profile by GPSC

| **GPSC** | **n** | **Predominant antimicrobial resistance profile**^a^ | **Continents** |
| --- | --- | --- | --- |
| 26 | 359 | CHL-SXT-TET (n=260) | Africa (n=261), Asia (n=62), Europe (n=25), North America (n=10), unknown (n=1) |
| 32 | 166 | pan-susceptible | Africa (n=4), Europe (n=6), North America (n=127), South America (n=29) |
| 55 | 101 | PEN (n=92) | Asia (n=91), Europe (n=1), North America (n=8), South America (n=1) |
| 334 | 101 | CHL-ERY-SXT-TET (n=35); ERY-SXT-TET (n=31) | Asia (n=97), Europe (n=3), Oceania (n=1) |
| 56 | 64 | pan-susceptible | Africa (n=64) |
| 212 | 12 | pan-susceptible | Europe (n=12) |
| 289 | 1 | SXT | South America |
| 575 | 1 | pan-susceptible | South America |
| 648 | 1 | SXT | Asia |

^a^The antimicrobial resistance profile includes antibiotic penicillin (PEN), chloramphenicol (CHL), erythromycin (ERY), cotrimoxazole (SXT) and tetracycline (TET).
