## Supplementary material for "Global population structure and phase variation of serotype 12F *Streptococcus pneumoniae* following the introduction of pneumococcal conjugate vaccine": Table S5

Table S5 Number of encapsulated and non-encapsulated serotype 12F strains

| Construct | Strain | Biological replicates | No. of encapsulated cells ± SD^a^ | No. of non-encapsulated cells ± SD | Encapsulated/non-encapsulated ratio |
| --- | --- | --- | --- | --- | --- |
| CPS12F | NUS0314 | 3 | 78 ± 55 | 1 ± 1 | 116.8 |
| Δ*cps* | NUS0114 | 3 | 0 | 180 ± 30 | 0 |
| Δ*wciJ*^12F^ | NUS7178 | 2 | 2 ± 3 | 596 ± 256 | 0 |
| *wciJ*^12F^ (195delA) | NUS7324 | 2 | 136 ± 118 | 33 ± 53 | 4.1 |
| *wciJ*^12F^ (195delA) backcrossed | NUS7389 | 2 | 67 ± 33 | 15 ± 7 | 4.4 |
